# Structure Determination of Inactive-State GPCRs with a Universal Nanobody

**DOI:** 10.1101/2021.11.02.466983

**Authors:** Michael J. Robertson, Makaía Papasergi-Scott, Feng He, Alpay B. Seven, Justin G. Meyerowitz, Ouliana Panova, Maria Claudia Peroto, Tao Che, Georgios Skiniotis

## Abstract

Cryogenic electron microscopy (cryo-EM) has widened the field of structure-based drug discovery by allowing for routine determination of membrane protein structures previously intractable. However, despite representing one of the largest classes of therapeutic targets, most inactive-state G protein-coupled receptors (GPCRs) have remained inaccessible for cryo-EM because their small size and membrane-embedded nature impedes projection alignment for high-resolution map reconstructions. Here we demonstrate that the same single-chain camelid antibody (nanobody) recognizing a grafted intracellular loop can be used to obtain cryo-EM structures of different inactive-state GPCRs at resolutions comparable or better than those obtained by X-ray crystallography. Using this approach, we obtained the structure of human neurotensin 1 receptor (NTSR1) bound to antagonist SR48692, of µ-opioid receptor (MOR) bound to the clinical antagonist alvimopan, as well as the structures of the previously uncharacterized somatostatin receptor 2 (SSTR2) in the apo state and histamine receptor 2 (H2R) bound to the H2 blocker famotidine. Each of these structures yields novel insights into ligand binding and specificity. We expect this rapid, straightforward approach to facilitate the broad structural exploration of GPCR inactive states without the need for extensive engineering and crystallization.

## Introduction

The impact of cryo-EM on structure-based drug discovery has been immense, leading to the characterization of a wide variety of new and valuable membrane protein drug targets, including ion channels and G protein-coupled receptors (GPCRs)^1^. As a key requirement for high-resolution cryo-EM map reconstructions, the randomly oriented particle projections must first be correctly aligned at intermediate-to-low resolution, a step that often fails for small proteins due to their limited structural features. For relatively small membrane proteins, the problem is compounded by the presence of a detergent micelle or lipid nanodisc used for solubilization, which further dampens the contrast of embedded protein densities. These obstacles pose a major bottleneck for cryo-EM structure determination of GCPRs, which account for roughly 35% of FDA-approved drugs. Approximately half of those are agonists that activate the receptor and the other half antagonists or inverse agonists^2^ that engage the inactive-state receptor, thereby blocking endogenous signaling. Active-state GPCR structures have been obtained by cryo-EM in complex with heterotrimeric G protein^3^, a ∼90 kDa entity that adds significant mass outside the detergent micelle or lipid disc and provides a source of alignment for the transmembrane receptor. However, there is no such generalizable tool for cryo-EM structure determination of inactive-state GPCRs, especially for family A receptors, the largest GCPR subfamily with hundreds of receptors that are typically ∼37-50 kDa and lack sizeable extracellular domains for projection alignment. Thus, the application of cryo-EM is severely limited for inactive-state GPCRs, and their structural elucidation mostly relies on extensive engineering and crystallization trials for X-ray studies. This limitation impacts mechanistic investigations of GPCRs and structure-based drug discovery for the countless therapeutically relevant receptors.

Che *et al*.^4^ recently reported a camelid VHH domain antibody, referred to as nanobody 6 (Nb6), that engages the inactive-state intracellular loop 3 (ICL3) of the κ-opioid receptor (KOR), even when this loop is grafted onto a wide range of receptors, including orphan receptors and family B receptors. This tool was initially developed as a bioluminescence resonance energy transfer (BRET)-based sensor of receptor activation state, an approach that also allows for cost-effective and robust screening of Nb6-binding constructs by employing BRET assays. Using the Nb6 approach, we obtained three-dimensional (3D) map reconstructions of human NTSR1 (hNTSR1) in complex with Nb6 bound to the inverse agonist SR48692^5^ at a resolution of 2.4 Å; the mouse MOR bound to a megabody^6^, an enlarged nanobody construct derived from Nb6 (Mb6), in complex with the antagonist alvimopan^7^ at a resolution of 2.8 Å; and the unliganded (apo) structure of human SSTR2 bound to Nb6 at 3.1 Å resolution. Further, a modified Nb6 (Nb6M) was cloned by grafting antigen-binding loops onto an alpaca nanobody scaffold and coupled with a nanobody-binding antibody fragment (NabFab^8^) to determine the structure of histamine receptor 2 (H2R) bound to famotidine at 3.0 Å resolution. Through these four structures, we demonstrate that Nb6 is a general solution for determining high-resolution inactive-state GPCRs by cryo-EM. Besides alleviating the need for extensive construct engineering and crystallogenesis, the cryo-EM/Nb6 approach has two distinct advantages: first, it does not necessitate the screening and development of receptor-specific nanobodies. Second, the Nb6 BRET-based sensor allows for rapid and efficient screening of constructs and conditions instead of time-consuming and expensive cryo-EM-based screening.

### Employing Nb6 for cryo-EM structure determination of GPCRs

Any fiducial marker used for cryo-EM structure determination of a target macromolecule needs to be rigid in its binding and orientation in respect to the structural target. Furthermore, if the marker is particularly small related to the target, it must be positioned to provide a distinct feature from various angles to allow unambiguous alignment of the randomly-oriented cryo-EM projections. A crystal structure of Nb6 bound to KOR revealed an interaction at the base of TM5 and TM6, off-center from the major axis of the receptor^4^. We hypothesized that this type of interaction would be an ideal asymmetric fiduciary for cryo-EM particle projection alignment as it would allow for the determination of the rotational orientation of the receptor around its major axis, particularly during initial low-resolution alignments.

To evaluate the degree of rigidity of Nb6 when bound to a GPCR, we first performed all-atom molecular dynamics (MD) simulations using the crystal structure of the KOR-Nb6 complex. Alignment of the MD trajectories on the receptor atoms and calculation of the average root mean squared fluctuation (RMSF) for Nb6 atoms over the final 100 ns of triplicate 500 ns simulations (Fig. 1a) showed that Nb6 is rigidly bound and characterized by little flexibility, except for the distal loops at the base of the nanobody.

**Fig 1.**
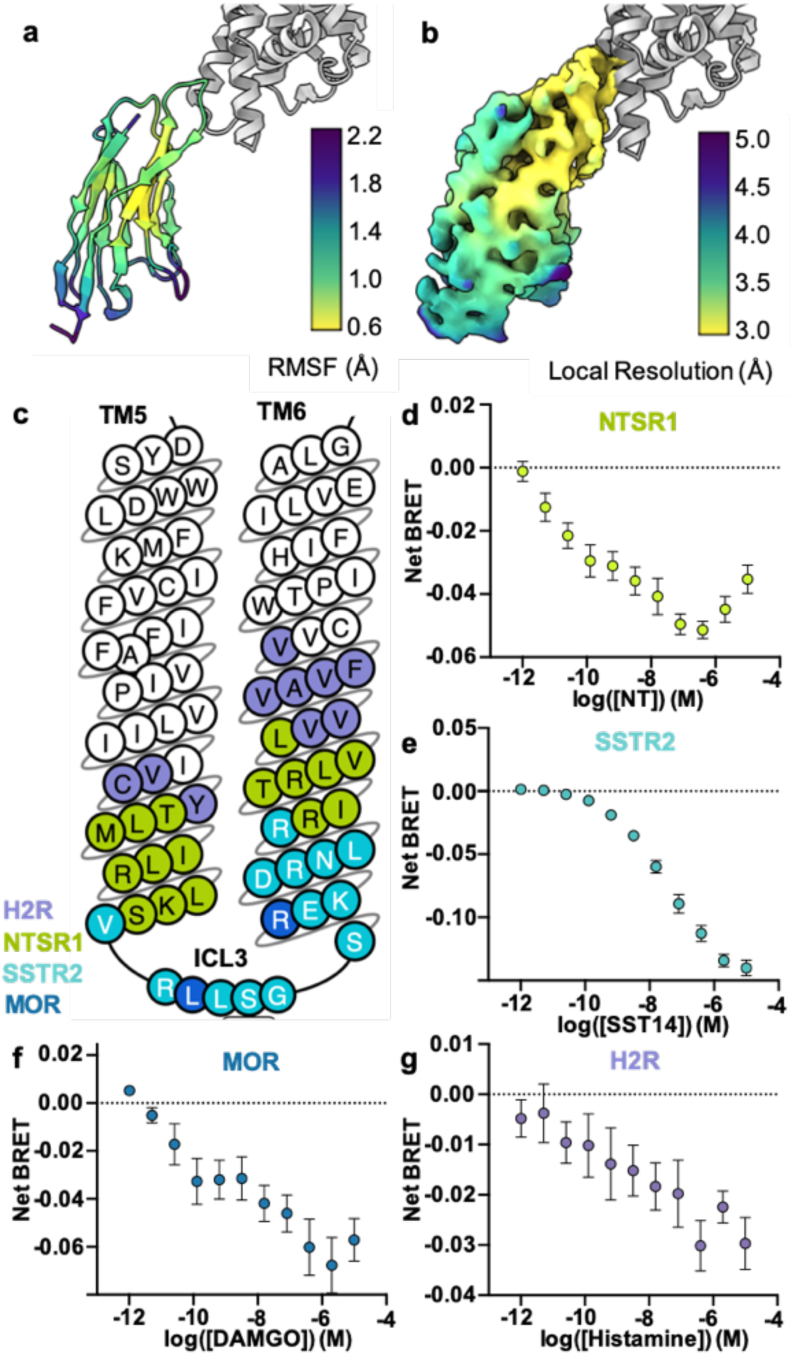
Construct design & evaluation for inactive state GPCR structure determination. **a**, Root mean squared fluctuations (RMSF) in atomic positions overaged over triplicate molecular dynamics simulations for Nb6 bound to KOR. **b**, local resolution plot of Nb6 from the SSTR2/Nb6 complex cryoEM map. **c**, overlay of the three KOR ICL3 constructs used to enable Nb6 binding for MOR (two point mutations), SSTR2 (5.68 to 6.31), and hNTSR1 (5.59 to 6.38). Dose-response curves representing loss of BRET transfer (net BRET) between Receptor-rLuc and Nb6-mVenus for **d**, NTSR1_κ_ and neurotensin (NT), **e**, MOR_κ_ and DAMGO, **f**, SSTR2_κ_ and SST14, and **g**, H2R_κ_ and histamine.

Encouraged by these computational predictions, we employed Nb6 for *de novo* cryo-EM structure determination of four pharmacologically relevant family A GPCRs. The first, NTSR1, is broadly associated with neurological and gastrointestinal processes^9^. Overexpression of NTSR1 in some tumor types promotes cancer progression and metastasis and is thus associated with poor prognosis^10^, prompting the study and development of NTSR1 inverse agonists for oncology^11^. We also studied MOR, the target of opioids, which are both the most effective treatment for pain and also the therapeutic class underlying the ongoing opioid epidemic in the United States^12^. While current opioid antagonists can treat acute opioid overdose, there is an urgent need for MOR antagonists with improved pharmacokinetic properties^12,13^. SSTR2 plays a pivotal role in the neuroendocrine system by opposing the release of many hormones, including growth hormone^14^. SSTR agonists are commonly prescribed for neuroendocrine tumors, while SSTR subtype-selective antagonists are in development for diabetes, but the lack of structural information for any somatostatin receptor family member in either active or inactive state has made the development of next-generation SSTR ligands challenging. Finally, selective H2R antagonists have a long history of clinical usage for peptic ulcers and gastroesophageal reflux disease and are amongst the most prescribed drugs. However, only structures of histamine receptor 1, the target for commonly prescribed drugs for allergic rhinitis and other allergies, have been determined to date, impeding a full understanding of histamine receptor subtype selectivity.

As originally described in Che *et al*., we engineered constructs to enable binding of Nb6 to mouse MOR and human SSTR2 by identifying minimal swaps of ICL3 along with the cytoplasmic tips of transmembrane helix 5 and 6 (TM5/6) with that of the KOR (Fig. 1c). In the case of the closely related MOR, the two point mutations M264L and K269^6.24^R (Ballesteros-Weinstein notation^15^; Fig. 1c) were sufficient to enable Nb6 binding. SSTR2 was able to bind Nb6 with the swap of all 15 residues from S238^5.68^ of TM5 (V256^5.68^ of KOR) to K252^6.31^ of TM6 (R270^6.31^ of KOR) (Fig. 1c). For human NTSR1 (hNTSR1) we utilized the same swap from KOR T247^5.59^ to L277^6.38^ as previously described for rat NTSR1^4^ (rNTSR1; Fig. 1c). Similarly, for human H2R we used an identical hybrid as described in Che *et al*. (KOR V244^5.56^ to V285^6.46^, Fig 1c). Notably, BRET assays of agonist-induced Nb6 dissociation confirmed construct functionality, with all four receptors showing dissociation of the nanobody in response to agonist binding (Fig. 1d-g). In all experimental replicates, the BRET data for MOR response to DAMGO appears to be biphasic, consistent with prior nanobody-based MOR activation sensors^16^, where distinct plasma membrane and endosomal signaling results in a two-phase response.

High-resolution cryo-EM structure determination was feasible in all cases. Despite Nb6 being only 12 kDa, Nb6, or the equivalent region of Mb6, is resolved in three maps, providing a critical stable density outside the detergent micelle surrounding NSTR1, MOR and SSTR2. Notably, the local resolution indications for Nb6 in our GPCR/Nb6 complex cryo-EM maps showed a high correlation with the relative root mean square fluctuation (RMSF) values for Nb6 calculated from our MD simulations (Fig. 1c). This observation further demonstrates the strong predictive power of MD simulations in assessing the behavior of fiducial markers for such work. In the case of H2R/Nb6M/NabFab, the entire complex is fairly rigid and well resolved, due, in part, to the nanobody modifications and/or the binding of the NabFab rigidifying the loops of Nb6M.

### Structure of hNTSR1

Obtaining inactive-state crystal structures has proven challenging for NTSR1. The only available crystal structures were determined in 2021^17^: one with a receptor in the apo state (3.19 Å) and several structures with the receptor bound to the inverse agonists SR48692 (2.64 Å, 2.70 Å) or SR142948A (2.92 Å), which are small molecules previously in development for treating cancer. Obtaining these structures required extensive engineering to facilitate protein crystallization, with that work employing a construct of rat NTSR1 evolved in bacteria to acquire numerous thermostabilizing mutations and fused in TM7 with a DARPin^18^ protein domain to facilitate crystal packing (rNTSR1-H4), although a structure was also obtained with a back-mutated construct (rNTSR1-H4bmx) at several positions to attempt to visualize a more native state^19^. To assess and compare the cryo-EM/Nb6 approach with crystallography we obtained the cryo-EM structure of hNTSR1/Nb6 complex bound to SR48692 (Fig. 2a, Extended Data Fig. 2). Notably, the cryo-EM structure was resolved at a global resolution of 2.4 Å, both a higher nominal resolution than the crystal structure and with better-resolved features, including the density corresponding to the ligand (Fig. 2a-e, Extended Data Fig. 3a, 3b, 8a). This resolution range also reveals densities corresponding to extensive hydration throughout the receptor in the cryo-EM map (Fig. 2f, Extended Data Fig. 3c).

**Fig 2.**
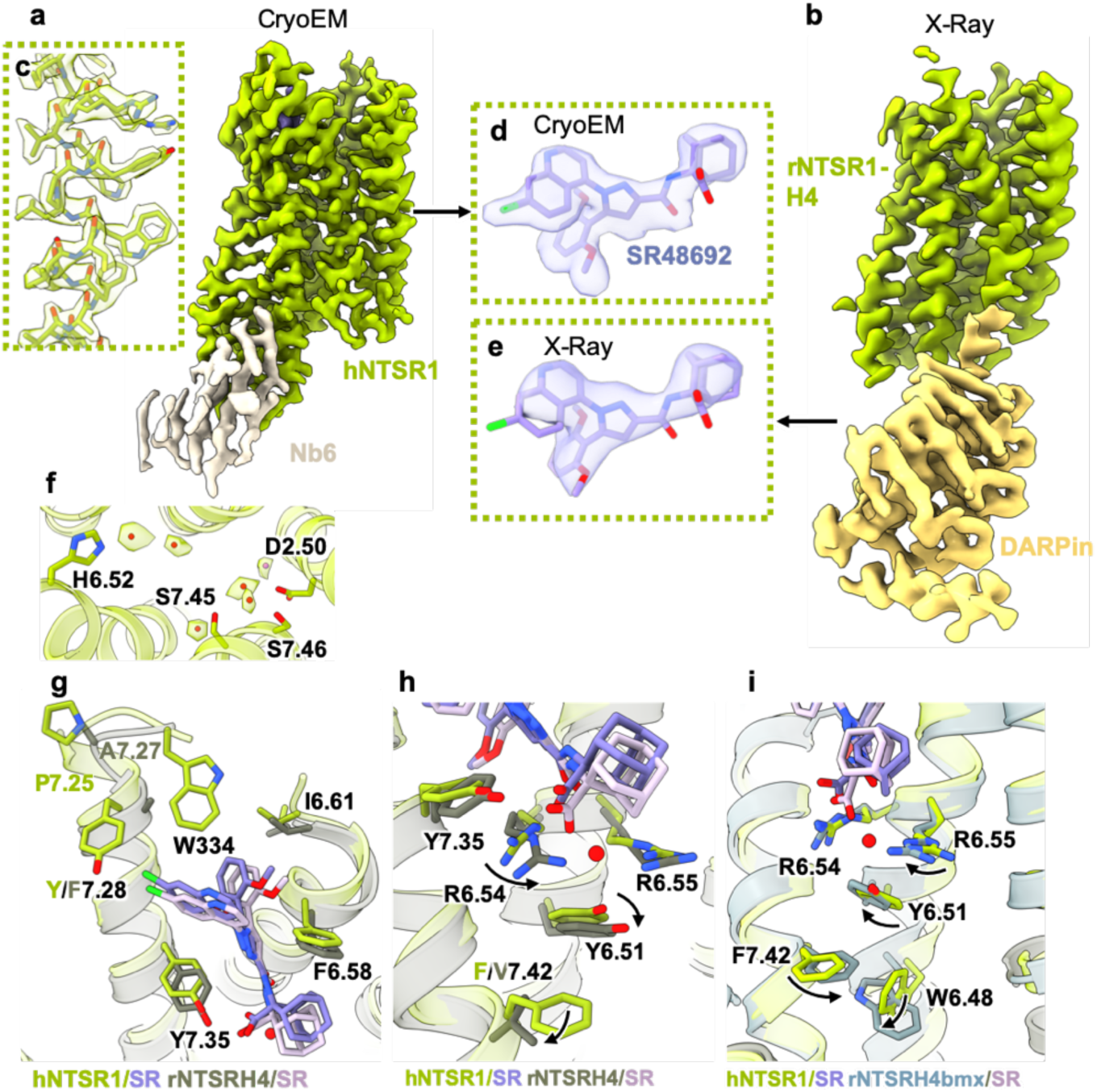
Comparison of cryoEM structure of hNTSR1 and crystal structure of rNTSR1-H4 and rNTSR1-H4bmx. **a**, 2.4 Å global resolution cryoEM map of hNTSR1. **b**, 2Fo-Fc crystallography map at 2.6 Å of rNTSR-H4 contoured at *σ*=1.0. **c**, hNTSR1 TM5 with cryoEM map **d**, Local EM map around bound inverse agonist SR48692. **e**, Local 2Fo-Fc map around SR48692 contoured at *σ*=1.25. **f**, Waters resolved in the core of the hNTSR1 receptor overlaid with the cryoEM map. **g**, Overlay of the hNTSR cryoEM structure (protein in green, SR48692 in purple) with the rNSTR-H4 crystal structure (protein in gray, SR48692 in pink) highlighting differences in ECL2. **h**, Overlay of the hNTSR cryoEM structure (protein in green, SR48692 in purple) with the rNSTR-H4 crystal structure (protein in gray, SR48692 in pink) highlighting the residue shift and probable loss of bound water molecule caused by the F7.42V thermostabilizing mutation **i**, Overlay of the hNTSR cryoEM structure (protein in green, SR48692 in purple) with the rNSTR-H4bmx crystal structure (protein in gray, SR48692 in pink)

The construct for hNTSR1 was minimally modified beyond the KOR ICL3 swap, with only N- and C-termini truncations in regions not resolved in typical family A structures and a single A85^1.54^L mutation to increase expression^20^. While the general ligand pose is very similar, the cryo-EM structure reveals additional interactions not observed in the rNTSR-H4 and/or rNTSR-H4bmx crystal structures. First, remodeling of the TM7-ECL3 region allows W334 in ECL3 to be fully resolved in the hNTSR1 structure loosely capping the top of the hydrophobic chloro-naphthyl and dimethoxy-phenyl moieties of SR48692 (Fig. 2g). This change may be due to several amino acid sequence differences at the top of TM7, most prominently P336^7.25^ in hNTSR1 versus T341^7.25^ in rNTSR1, with the proline inducing a sharp turn between the top of TM7 and ECL3 in hNTSR1. In addition, a rearrangement of R327^6.54^ in rNTSR-H4 (R322^6.54^ in hNTSR1) appears to disrupt a water-mediated interaction between R327^6.54^, R328^6.55^ (R323^6.55^ in hNTSR1), and the carboxylate moiety of SR48692 observed in the hNTSR1 structure (Fig. 2h). This R327^6.54^ rearrangement likely stems from the F358^7.42^V thermostabilizing mutation in rNTSR-H4, allowing a downward relaxation of Y324^6.51^ (Y319^6.51^ in hNTSR1) and R327^6.54^. This water molecule is also displaced in the rNTSR-H4bmx structure due to a different rearrangement of R327^6.54^ and R328^6.55^, likely resulting from a shift in the W321^6.48^ rotamer not observed in either hNTSR1 or rNTSR-H4bmx (Fig 2i). Notable differences also exist in the TM helices between the two structures; TM1 is shifted with respect to the core of the receptor in both crystal structures in varying degrees and directions (Extended Data Fig. 3d). In addition, the top of TM2 and ECL1 are out of register by one residue between hNTSR1 and rNTSR-H4. While this may be due to the differences in experimental constructs, the observed discrepancy may also result from the fact that even at α=1 the 2Fo-Fc map of the crystal structure lacks sidechain information in the ECL1 region, rendering modeling ambiguous (Extended Data Fig. 3e,f). Overall, the lack of thermostabilizing mutations and DARPin in TM7 restores a more native-like inactive state as suggested by the conformation of key amino acids. For example, Y364^7.53^ of the hNTSR1 NPxxY motif (Y369^7.53^ in rNSTR1) occupies a position and rotamer more typical of an inactive state family A GPCR (Extended Data Fig. 3g).

### Structure of MOR

Using the same approach, we obtained a cryo-EM structure of the MOR bound to the non-blood brain barrier penetrant, MOR selective inverse agonist alvimopan, an FDA-approved drug for reversing opioid-induced gastrointestinal (GI) symptoms in hospital settings. In the case of NTSR1/Nb6, vitrifying the sample in a very thin buffer layer on holey gold grids^21^ was necessary for high resolution and to achieve proper alignment of particle projections. To assess an alternative approach, we used Nb6 to develop a megabody, termed ‘Mb6’, a fusion of a nanobody and a scaffold protein of either 45 or 86 kDa that can serve as a larger fiducial marker, based on the work of Uchański *et al*.^6^. In that work, the Mb designs employed for GABA_A_ and WbaP resulted in cryo-EM maps that could not resolve well past the nanobody portion, presumably due to flexibility. We thus opted instead for using the c7HopQA12 design, which incorporates a twisted linker between the nanobody and the 45 kDa HopQ scaffold to increase their contact and, ideally, rigidity. However, our MD simulations suggested that even this design was characterized by substantial flexibility between the Nb and scaffold (Extended Data Fig. 5a). In agreement with this result, our cryo-EM 2D class averages of the MOR-Mb6 complex showed that only the Nb6-receptor portion is well resolved, with the scaffold averaging out (Extended Data Fig. 4b). Nevertheless, leveraging the ordered Nb6 portion of Mb6 in our cryo-EM data, we obtained a 2.8 Å cryo-EM map (Fig. 3a, Extended Data Fig. 4) with unambiguous density for the ligand and receptor (Fig. 3c, Extended Data Fig. 8b). This level of detail is quite comparable to the 2.8 Å crystal structure of MOR (Fig. 3b, d), although the cryo-EM map further resolves several probable water molecules in the ligand binding site, including one that bridges interactions between the carboxylate of alvimopan and the receptor (Extended Data Fig. 5b).

**Fig 3.**
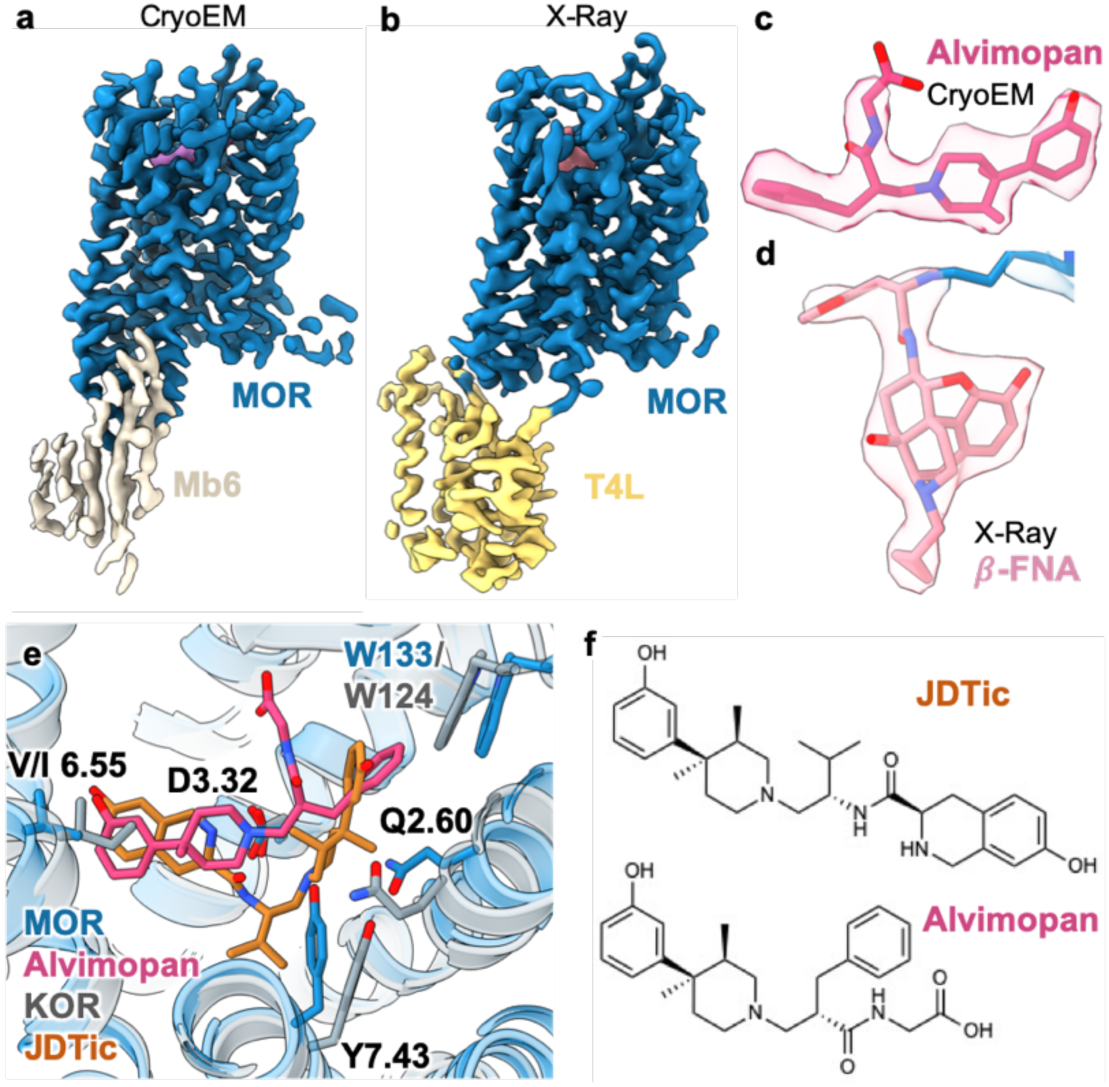
Comparison of cryo-EM structure of alvimopan-bound MOR with crystal structures of MOR/*β*-FNA and KOR/JDTic. **a**, Cryo-EM map of MOR at 2.8 Å global resolution. **b**, 2Fo-Fc crystallography map of MOR at 2.8 Å contoured at *σ*=2.0. **c**, Cryo-EM density and model for inverse agonist alvimopan. **d**, Local 2Fo-Fc map around *β*-FNA contoured at *σ*=1.5. **e**, overlay of the alvimopan pose in the MOR cryo-EM structure (protein in blue, alvimopan in magenta) with the JDTic pose in the KOR crystal structure (protein in grey, JDTic in orange). **f**, Comparison of JDTic and alvimopan chemical structures highlighting similar phenol-piperidine scaffold.

Comparing the binding poses of the MOR-selective alvimopan with that of the structurally similar ligand JDTic, which is selective for KOR, reveals a striking difference. Despite their similar scaffolds, the phenol-piperidine moieties of alvimopan and JDTic are in completely different positions and orientations, nearly flipped in the binding pocket. Aligning our structure with KOR-JDTic (PDB:4DJH^22^) reveals that JDTic would clash with Y326^7.43^ and Q124^2.60^ in MOR, while alvimopan would clash with W124 and I294^6.55^ in KOR (I294^6.55^ of KOR is substituted by a repositioned valine in MOR). In addition to the steric confines of the binding pocket, charge differences also appear to contribute to the differences in binding pose between JDTIC and alvimopan. The nitrogen of the tetrahydroisoquinoline ring system of JDTic is likely also protonated or perhaps protonated instead of the piperidine, and thus forms salt bridge interactions with D138^3.32^ (D147^3.32^ in MOR), just as the protonated piperidine of alvimopan does in MOR. Of note, alvimopan shares similarities to fentanyl and its analogues, including a central piperidine ring. Computational docking of carfentanil^23^, a potent MOR agonist, in the active-state structure of the receptor^3^ suggests a pose similar to alvimopan, but with the additional phenylamine moiety on the piperidine ring positioned to bind into the pocket occupied by the toggle switch tryptophan W293^6.48^ (Extended Data Fig. 5a). How a MOR ligand engages this region is often the main determining factor for its agonism versus antagonism. Based on our observations, we hypothesize that this phenyl moiety of fentanyl and its derivatives is responsible for converting the piperidine scaffold to an agonist.

### Structure of SSTR2

To test the cryo-EM/Nb6 approach on a receptor without prior structural characterization we chose SSTR2, a member of the SSTR subfamily for which no structures have been determined. Despite the absence of a stabilizing inverse agonist or any other ligand, and not using an energy filter for cryo-EM data acquisition (typically beneficial for improved contrast when imaging small proteins), we were able to obtain a 3.1 Å map of SSTR2/Nb6 with well-resolved features (Fig. 4a, Extended Data Fig. 6, 8c). Together with two active-state structures of SSTR2 bound to two different agonists, the endogenous 14-mer SST14 and the synthetic 8-mer octreotide^24^, the inactive-state SSTR2 structure enabled us to characterize several aspects of receptor activation and ligand recognition. Perhaps the most interesting feature of SSTR2 is the conformational flexibility of ECL2; while in a well-defined position in the apo state structure, ECL2 is pushed to a more open conformation when bound to the endogenous agonist SST14, but folds down over the top of the receptor in the octreotide-bound state (Fig. 4b). As a consequence of the ECL2 plasticity, different residues are engaged in binding the pan-SSTR agonist SST14 versus the SSTR2-selective octreotide. This is an important factor to SSTR subtype selectivity, which is explored further in our recent study^24^.

**Fig 4.**
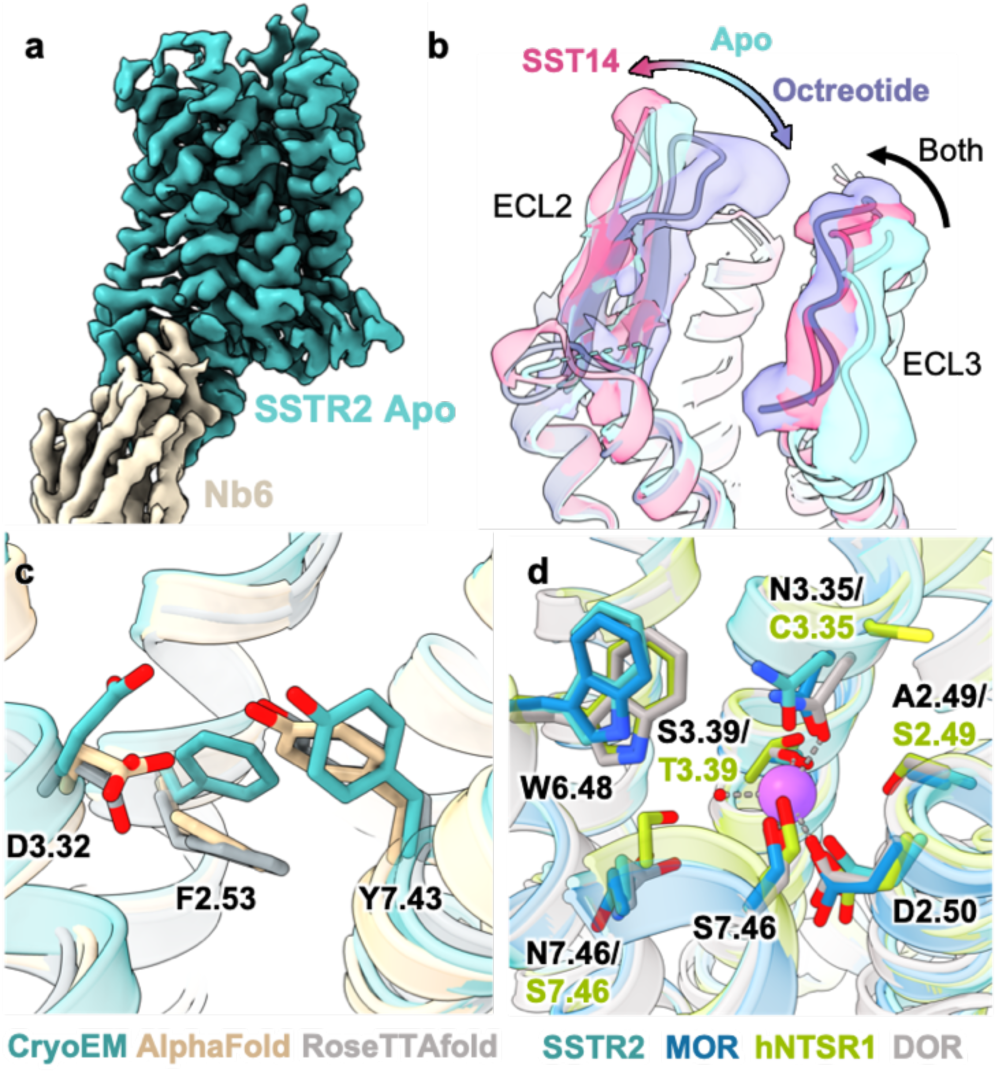
Cryo-EM structure of the apo SSTR2 and comparison of sodium ion binding site. **a**, Cryo-EM map of SSTR2 at 3.1 Å global resolution. **b**, Comparison of the model and cryo-EM maps of apo SSTR2 (teal) and SSTR2 in complex with either SST14 (magenta) or octreotide (purple). **c**, Comparison of the CryoEM structure of SSTR2 (teal) with AlphaFold (tan) and RoseTTAfold (gray) predictions. **d**, Overlay of SSTR2 (teal), MOR (blue), and NTSR1 (green) cryo-EM structures around the canonical family A sodium ion binding site with the DOR sodium coordination site structure (gray, PDB:4N6H).

Very few structures have been obtained to date for family A GPCRs in the unliganded, apo state, primarily due to the general need for a stabilizing ligand to obtain a crystal structure. Experimentally-derived apo structures provide an opportunity to evaluate predicted structures from the recently published deep-learning based tools, AlphaFold2^25^ and RoseTTAfold^26^. Notably, while both software performed well in predicting the overall structure of SSTR2, key details of the ligand binding site were in disagreement with the experimental structure, particularly in residues D122^3.32^, F92^2.53^, and Y302^7.43^ (Fig. 4c), which are involved in ligand binding and activation, as detailed in the companion active-state SSTR2 manuscript^24^. Although a more systematic comparison is needed, this observation underscores both the merits and the potential limitations of homology model-based approaches. Differences in side chain positioning, especially within a ligand binding site, can throw off structure-based drug discovery efforts and other pursuits requiring high-accuracy coordinates.

### Differential sodium accommodation by family A GPCRs

Our near-native inactive-state cryo-EM maps provide the opportunity to examine the allosteric sodium binding site found deep in the core of many family A receptors. For NTSR1, MOR, and SSTR2 there is strong experimental evidence for negative allosteric modulation by sodium^27–29^. In both MOR and SSTR2, the identities of the residues in the sodium binding site are the same as the δ-opioid receptor (DOR), for which there is a high-resolution crystal structure (1.8 Å) with a well-defined sodium site^30^ (Fig. 4c). The geometries of the MOR, SSTR2, and DOR sites are very similar, with the exception of a slight rearrangement of W^6.48^ in the case of both MOR (W293^6.48^) and SSTR2 (W269^6.48^), and N125^3.35^ in SSTR2. In contrast to SSTR2 and MOR, hNTSR1 has four amino acid differences in the sodium binding site, namely T155^3.39^ (in place of S^3.39^), S357^7.46^ (in place of N^7.46^), S111^2.49^ (in place of A^2.49^), and C151^3.35^ (in place of N^3.35^). In the DOR structure, both S135^3.39^ and N131^3.35^ directly interact with the sodium.

The three maps we determined resolve features consistent with water and/or ions in the canonical sodium binding region near D^2.50^, although only NTSR1 has sufficiently high resolution in this region to facilitate unambiguous ion modeling (Extended Data Fig. 7a-c). The probable ion sites of SSTR2 and MOR match relatively well to those observed in DOR. In the hNTSR1 map, we observe a density with close proximity to oxygen atoms of T155^3.39^, S111^2.49^, D112^2.50^ (Extended Data Fig. 7a). Even though precise identification of water versus ion is challenging in cryo-EM maps, we expect that this feature corresponds to the sodium, shifted by 1.8 Å compared to the sodium in DOR (Fig. 4d, Extended Data Fig. 7d, e). As other receptors with resolved sodium ion sites have cysteine or other hydrophobic residues in place of N^3.35^, S111^2.49^ is likely responsible for the shift in sodium position in NTSR1. Overall, we find substantial similarity between the sodium binding sites of inactive state GPCRs from cryo-EM and crystallography.

### Structural Basis of Histamine Receptor Subtype Selectivity

Notwithstanding the success of Nb6 alone to obtain these structures, we sought to generate a larger, rigid Nb6-based fiducial that would enable the alignment of potentially more challenging targets and also extend this approach to less powerful electron microscopy platforms. To this end we generated an ‘alpacaized’ Nb6 where the antigen binding loops of Nb6 were grafted onto an alpaca nanobody scaffold^8^. This modified Nb6, called Nb6M, was used to bind H2R possessing the KOR ICL3 swap in complex with the FDA approved H2 blocker famotidine. The complex was additionally decorated with a NabFab^8^ (a Fab antibody fragment that binds alpaca nanobodies) and an anti-Fab nanobody that has been shown to reduce Fab flexibility^31^. The structure of the entire complex was determined by cryo-EM at a global nominal resolution of 3.0 Å (Fig. 5a, Extended Data Fig. 8). With the exception of the anti-Fab nanobody, the structure is well resolved, with local refinement on the receptor and Nb6M region further enhancing the details of H2R and bound famotidine (Fig. 5b, Extended Data Fig. 8e). Notably, a lipid molecule can be observed wedged between TM1 and TM7 and interacting with Q79^2.62^, in a similar position observed previously in cryoEM maps with certain family A GPCRs^32^ although its role remains unclear (Extended Data Fig. 9a).

**Fig 5.**
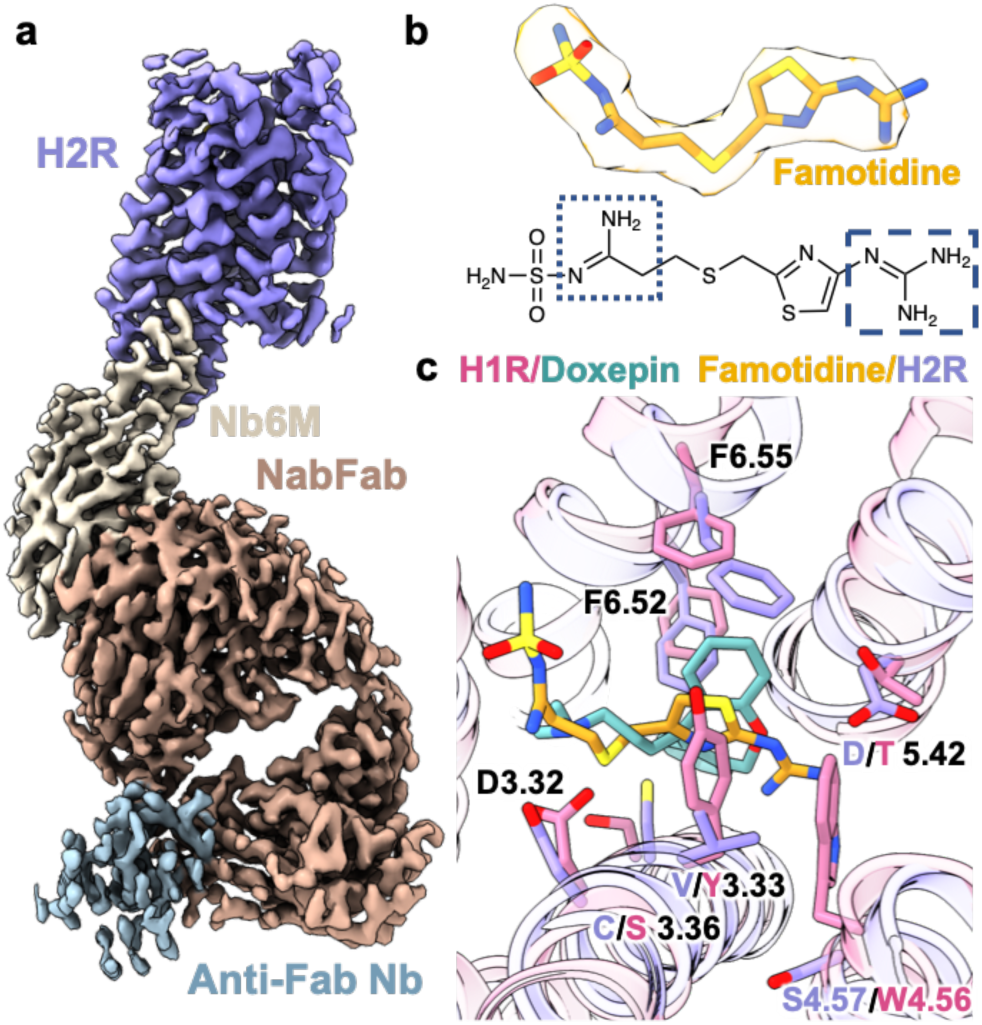
Cryo-EM structure of the H2R/Famotidine/Nb6M/NabFab/A nti-Fab complex and comparison to H1R/Doxepin. **a**, 3.0 Å global resolution cryo-EM map of inactive-state H2R complex. **b**, Famotidine in the cryo-EM density and chemical schematic; protonatable amine groups are boxed in blue, with the moiety protonated near neutral pH on the right in large dashed lines, and the moiety likely protonated at low pH on the left in small dotted lines. **c**, Overlay of H1R (magenta) and doxepin (teal) crystal structure with the H2R (lavender) and famotidine (gold) cryo-EM structure.

Comparing our cryo-EM structure of H2R/famotidine with the crystal structure of H1R bound to the H1R selective doxepin (PDB:3RZE^33^) reveals clearly the structural basis for subtype selectivity between H1R and H2R. H1R ligands, including the FDA approved doxepin, loratadine, and cetirizine (Extended Data Fig. 9b-d) possess a distinct diphenyl group that occupies a large hydrophobic pocket in H1R comprised of W^4.56^, F^6.55^, F^6.52^, and Y^3.33^ (Fig. 5c). These compounds also include a single amine which is protonated at neutral pH and interacts with the key aminergic aspartate D^3.32^ present in many family A GPCRs. The hydrophobic pocket where the diphenyl moiety binds in H1R is substantially perturbed in H2R, with the substitution of Y^3.33^V and repositioning of W^4.56^ in H1R with S^4.57^ in H2R removing major hydrophobic and pi-pi interaction partners (Fig. 5c). Further, TM6 shifts inward, repositioning F^6.55^ and F^6.52^ and resulting in a partial closure of the pocket (Fig. 5c). Several additional residues, including C^3.36^ (S^3.36^ in H1R) occupy rotamers in the famotidine-bound state that would sterically clash with doxepin binding.

Famotidine and doxepin differ in their chemical structures, where famotidine is a primarily linear molecule with two amine-containing moieties (Fig. 5b), a common feature of H2R blockers (Extended Data Fig. 9e-g). These two amine-containing groups of H2R antagonists tend to differ substantially in their pKas, with one that is likely protonated at or near pH 7.0 (the guanidine group in the case of famotidine, Fig. 5b; dashed lines) and a second that would require a far lower pH for protonation (the amidine/thiazole region for famotidine^34^, Fig. 5b, dotted lines). However, as H2R antagonists are primarily used for diseases of gastric acid and bind to hydrochloric acid-secreting parietal cells, the precise protonation state in the therapeutic environment is difficult to predict. Regardless of protonation state, the binding pose is likely very similar; the protonated high-pKa amine moiety forms a salt bridge with D^5.42^ while the other amine-containing group interacts with D^3.32^ via either hydrogen bonding or a salt bridge (Fig. 5c). Prior functional studies have also suggested that D^5.42^ is a main determinant in subtype selectivity for H2R blockers^35^, with this position corresponding to a threonine in H1R, which is unable to form the key salt bridge. Our docking studies with the other FDA approved H2R blockers cimetidine and ranitidine suggest this ‘di-aminergic’ ligand binding at H2R is a common feature of H2R antagonists (Extended Data Fig. 9e-g). Additional factors contributing to the lack of binding of famotidine to H1R include the steric clash with W^4.54^ (S in H2R) and the modulation of the shape of the hydrophobic binding pocket (Fig. 5c).

## Discussion

Here, we present inactive-state structures of four family A GPCRs, all obtained by cryo-EM at resolutions as high as 2.4 Å by employing minimal engineering and a universal nanobody for projection alignment. By using molecular dynamics simulations to assess fiducial markers for use in cryo-EM structure determination, we could accurately predict that Nb6 is sufficiently rigid for cryo-EM particle alignment, while Mb6 is too flexible for resolution beyond the nanobody portion. Additionally, the enhanced decoration opportunities afforded by the alpacaized Nb6M provide an additional tool for structure determination of potentially more challenging targets and with less powerful cryo-EM imaging set-ups such as those of 200kV and even 100kV. Each of the cryo-EM structures described here yields new insights into ligand recognition, with resolution that is as good or better than that of analogous structures produced by X-ray crystallography. In contrast to active state GPCR-G protein complexes, where the highest-resolution portion of the map is on the larger G protein due to the center of alignment, the inactive-state Nb6 GPCR structures display their highest resolution at the receptor core, facilitating accurate modeling of ligands and their surrounding hydration. Even with the inclusion of NabFab, local refinement on the receptor-Nb6M yielded a very well resolved ligand binding cavity. This generalizable approach opens this broad family of drug targets to rapid, crystallization-free structure determination, with biochemical stability of the receptor and nanobody binding being the only limitations. As such, we anticipate Nb6 to be a broadly applicable approach to determining inactive state structures of near-native GPCRs, further propelling drug discovery in this class of receptors.

## Supporting information

All supplemental files

## Author Contributions

M.J.R. cloned constructs, expressed and purified proteins, processed EM data, built models, and ran/analyzed molecular dynamics simulations. M.P.S. expressed and purified proteins and collected cryo-EM data. F.H. cloned constructs and expressed proteins. J.M. performed BRET assays. A.S. prepared cryo-EM samples and collected cryo-EM data. M.C. expressed proteins and purified nanobody. O.P. prepared cryo-EM samples and collected cryo-EM data. T.C. provided constructs for nanobody expression and BRET assays. M.J.R. and G.S. wrote the manuscript with input from M.P.S., F.H., J.M., O.P., and T.C.. G.S. supervised the project.

## Data Availability

All data generated or analyzed in this study are included in this article and the Supplementary Information. The cryo-EM density maps and corresponding coordinates have been deposited in the Electron Microscopy Data Bank (EMDB) and the Protein Data Bank (PDB), respectively, under the following accession codes: NTSR1 (EMD-26589, PDB:7UL2), H2R (EMD-26590, PDB:7UL3), MOR (EMD-26591, PDB:7UL4), SSTR2 (EMD-26592, PDB:7UL5). Raw data has been deposited in EMPIAR under the following accession codes:

## Acknowledgements

We thank the Kobilka lab for providing plasmids of NTSR1 and MOR. We thank the Kossiakoff lab for providing the plasmid for NabFab and constructive feedback. Cryo-EM data were collected at the Stanford cryo-EM center (cEMc) with support from E. Montabana. This work was supported, in part, by the Mathers Foundation (G.S.), training grant T32GM089626 (J.G.M.), the National Institute of Health R35GM143061 (T.C.), and used the Extreme Science and Engineering Discovery Environment (XSEDE)^36^ resource comet-gpu through sdsc-comet allocation TG-MCB190153 (G.S.), which is supported by National Science Foundation grant number ACI-1548562.

## Methods

### Cloning

The cDNA for SSTR2 was obtained from Horizon Discovery and cloned into a pFastBac vector containing an N-terminal haemagglutinin (HA) signal sequence followed by a FLAG epitope (DYKDDDD) and a C-terminal C3 protease cleavage site followed by enhanced green fluorescent protein (eGFP) and a hexahistidine (His6) tag using Gibson cloning. Constructs for hNTSR1 and MOR were obtained from the lab of Brian Kobilka and have been described previously^3,20^. Briefly, each is cloned in pFastBac with an N-terminal haemagglutinin (HA) signal sequence followed by a Flag epitope (DYKDDDD) and a C-terminal HRV 3C protease cleavage site followed by a hexahistidine (His6) tag. In the case of hNTSR1, the His6 tag is preceded by enhanced green fluorescent protein (eGFP). For MOR, two point mutations (M264L and K269R) were introduced with site-directed mutagenesis; all PCR reactions were conducted with Q5 polymerase.

For hNTSR1, residues between T247^5.59^ to L277^6.3^ of kappa opioid were substituted, for SSTR2, residues between S238 and K252 were replaced with those of kappa opioid receptor, and for H2R residues from V244^5.56^ to V285^6.46^ were exchanged for kappa opioid.

### Biolumescence resonance energy transfer (BRET) assays

BRET assays were performed and analyzed as previously described^4^ with the following modifications: HEK-293S cells grown in FreeStyle 293 suspension media (Thermo Fisher) were transfected at a density of 1 million cells/mL in 2 mL volume using 600 ng total DNA at 1:1 ratio of Receptor-rLuc:Nb6-mVenus and a DNA:PEI ratio of 1:5, and incubated in a 24 deep well plate at 220 rpm, 37°C for 48 hours. Cells were harvested by centrifugation, washed with Hank’s Balanced Salt Solution (HBSS) without Calcium/Magnesium (Gibco), and resuspended in assay buffer (HBSS with 20 mM HEPES pH 7.45) with 1 μg/mL freshly prepared coelenterazine h (Promega). Cells were plated in white-walled, white-bottom 96 well plates (Costar) in a volume of 60 μl/well and 60,000 cells/well. Ligands were prepared in drug buffer (assay buffer with 0.1% BSA, 6 mM CaCl_2_, 6 mM MgCl_2_), and added at a 1:2 ratio of drug:cell suspension. Ten minutes after the addition of ligand, plates were read using a SpectraMax iD5 plate reader using 485 nm and 535 nm emission filters with a one-second integration time per well. The computed BRET ratios (mVenus/RLuc emission) were normalized to ligand-free control (Net BRET) prior to further analysis.

### Expression and Purification of Nb6, Mb6, and Nb6M

Wk6 *E. coli* (ATCC) were transformed via heat shock with Nb6 plasmid^4^ and two 5 mL starter cultures were grown overnight in LB supplemented with 100 µg/ml ampicillin. 2L of terrific broth were supplemented with 100 µg/ml ampicillin, inoculated with starter cultures, and grown at 37°C with shaking until OD_600_=0.7. Expression was then induced with 1mM IPTG and expression was allowed to proceed at 28°C overnight. Pellets were harvested via centrifugation and washed once with phosphate-buffered saline before snap freezing in liquid nitrogen. Pellets containing Nb6 were thawed and resuspended in 50 mM Tris pH 8.0, 0.5 mM EDTA, 20% w/v sucrose (TES) buffer at 15 mL / 1 liter pellet and supplemented with protease inhibitor cocktail and shaken in an orbital shaker for 1 hour at 4°C. An additional 30 mL per 1 liter pellet of TES diluted 1:4 with water was added and placed in an orbital shaker for an additional 45 minutes at 4°C. Cell debris was removed by ultracentrifugation at 100,000xg for 30 minutes. The supernatant was filtered with a 45 micron filter, supplemented with 20 mM imidazole, and loaded over a gravity Ni-NTA column at 4°C. The column was washed with 10 CV of a buffer containing 250 mM NaCl, 50 mM Tris pH 7.5, 10 mM imidazole, and then eluted with buffer containing 250 mM imidazole. Nanobody was concentrated and applied to size exclusion chromatography with a buffer containing 250 mM NaCl and 50 mM Tris pH 7.5. Monomeric fractions were pooled, concentrated, supplemented with 10% glycerol, and snap frozen in liquid nitrogen for later use. Expression and purification of Mb6 and Nb6M was performed identically to Nb6.

### Expression and Purification of NabFab

Chemically competent C43 *E. coli* were transformed via heat shock with NabFab plasmid^8^. A culture of 10 mL 2xYT media (16g/L tryptone, 10g/L yeast extract, 5g/L NaCl) was inoculated with a single colony and incubated at 37°C with shaking overnight. TB autoinduction media (Terrific Broth supplemented with 0.4% glycerol, 0.01% glucose, 0.02% lactose,1.25 mM MgSO_4_, and 100 µg/ml ampicillin) was inoculated from starter cultures and incubated for 6 hrs at 37 °C, 225 rpm, before decreasing the temperature to 30°C for overnight growth. The cells expressing NabFab were harvested by centrifugation and snap-frozen in liquid nitrogen for storage. To purify NabFab, cell pellets were thawed and resuspended in lysis buffer containing 20 mM sodium phosphate, pH 7.4, 150 mM NaCl, DNaseI, and protease inhibitor cocktail), and lysed by sonication. The lysate was then incubated at 63-65 °C for 30 minutes before centrifugation at 20,000 xg for 30 minutes. The supernatant was collected and run over a Protein G column pre-equilibrated with 20 mM sodium phosphate, 500 mM NaCl, pH 7.4. The NabFab was then eluted with 0.1 M acetic acid and immediately loaded onto a Resource S cation exchange column pre-equilibrated with buffer A (50 mM sodium acetate, pH 5.0). The column was washed with 5 CV of buffer A and protein then eluted over a salt gradient 0-100% buffer B (50 mM sodium acetate, 2M NaCl, pH 5.0). Eluted NabFab was dialyzed overnight into 150 mM NaCl, 20 mM HEPES, pH 7.5 and then concentrated to 3 mg/mL for complexation.

### Expression and Purification of Receptor/Nb6 Complexes

All receptors were expressed in Sf9 insect cells (Expression Systems) infected at a density of 3-4 million cells/ml. At 48 hours post-infection, cells were collected with centrifugation, washed with phosphate-buffered saline containing 10 µM antagonist (when used), and pellets were snap frozen in liquid nitrogen for purification. A general protocol was used for producing receptor/Nb6 complexes (Fig. S1), with some slight deviations. Sf9 pellets containing receptor were lysed in hypotonic buffer containing 20 mM HEPES pH 7.5, 5 mM MgCl_2_, protease inhibitor cocktail, benzonase, 10 µM antagonist (when used), and 2 mg/ml iodoacetamide and gently stirred for an hour at 4°C. Membranes were harvested with ultracentrifugation at 100,000xg, supernatant was discarded, and membranes were resuspended in solubilization buffer containing 250 mM NaCl, 20 mM HEPES pH 7.5, 1 mM MgCl_2_, protease inhibitor cocktail, 10 µM antagonist (when used), and 2 mg/ml iodoacetamide; then drip frozen into liquid nitrogen and stored at -80°C. Resuspended membranes were thawed and detergent was added dropwise while stirring at 4°C to a final concentration of 1% LMNG/0.1% CHS/0.1% Cholate. After 3 hours, insoluble debris was removed with ultracentrifugation at 100,000 g. Solubilized receptor was supplemented with 20 mM imidazole and gravity loaded over a Ni-NTA resin column. Columns were washed with 10 column volumes of buffer containing 250 mM NaCl, 20 mM HEPES pH 7.5, 10 µM antagonist (when used), 20 mM imidazole, and 0.1% LMNG/0.01% CHS and protein was eluted in buffer containing 250 mM NaCl, 20 mM HEPES pH 7.5, 10 µM antagonist (when used), 250 mM imidazole, and 0.01% LMNG/0.001% CHS. Receptor was then supplemented with 5 mM CaCl_2_ and loaded onto M1 flag resin; washed with 5 column volumes of 250 mM NaCl, 20 mM HEPES pH 7.5, 10 µM antagonist (when used), 2 mM CaCl2, and 0.1% LMNG/0.01% CHS; and eluted with buffer containing 150 mM NaCl, 20 mM HEPES pH 7.5, 10 µM antagonist (when used), 1 mM EDTA, 0.2 mg/ml FLAG peptide, and 0.1% LMNG/0.01% CHS. Purified receptor was incubated on ice overnight with HRV 3C protease and purified Nb6 was added at a 2:1 molar ratio. Receptor/Nb6 complex was concentrated in a 50 kDa molecular weight cutoff spin concentrator and subjected to SEC chromatography with an ENrich 650 column (Bio-Rad) and a buffer containing 150 mM NaCl, 20 mM HEPES pH 7.5, 0.001% LMNG, 0.00033% GDN, and 0.0001% CHS. Fractions containing monomeric receptor/Nb6 complex were pooled for cryo-EM sample preparation. This protocol was identical for SSTR2, NTSR1, and H2R with the exception of the addition of 100 µM TCEP to the lysis buffer for NTSR1 and incubation with Nb6M, NabFab, and anti-Fab Nb (Thermo Fisher Scientific 1033270500) in the case of H2R. Preparation of MOR/Mb6 complex was identical to that of receptor Nb6 complex, with the omission of the HRV-3C cleavage to remove eGFP as it was not present in the construct.

### Cryo-EM Sample Preparation

All samples were prepared on glow-discharged holey gold grids (Quantifoil ultrAufoil R1.2/1.3), blotted in a FEI Vitrobot Mark IV (Thermo Fisher Scientific) at 4°C and 100% humidity, and plunge frozen into liquid ethane. Blotting conditions for each sample were as follows: 3.0 µl of NTSR1/Nb6 complex at 16 mg/ml with an additional 0.05% beta OG; 3.0 µl of MOR/Mb6 complex at 5 mg/ml; 2.5 µl of SSTR2/Nb6 complex at 16 mg/ml with an additional 0.05% beta OG; and 3.0 µl of H2R/Nb6M/NabFab/anti-Fab Nb at 2.5-4 mg/ml.

### Cryo-EM Data Collection

All samples were collected on a KRIOS electron microscope at an accelerating voltage of 300 kV, with an energy filter for NTSR1/Nb6, H2R/Nb6M/NabFab/anti-Fab Nb, and MOR/Mb6 and without an energy filter for SSTR2/Nb6. All data was collected through SerialEM with beam tilt compensation and recorded on a Gatan K3 direct electron detector. The resulting image stacks have a pixel size of 0.434 Å for NTSR1/Nb6, H2R/Nb6/NabFab, MOR/Mb6, and H2R/Nb6M/NabFab/anti-Fab Nb; and 0.426 Å for SSTR2/Nb6, all in super resolution mode. Each NTSR1/Nb6 image stack is composed of 71 frames with an incident electron dose of 0.85 e^-^/Å^2^ per frame, for a total dose of 61 e^-^ /Å^2^/s per micrograph. Each MOR/Mb6 image stack is composed of 63 frames with an incident electron dose of 0.98 e^-^/Å^2^ per frame, for a total dose of 61 e^-^/Å^2^/s per micrograph. Each SSTR2/Nb6 image stack is composed of 55 frames with an incident electron dose of 1.26 e^-^/Å^2^ per frame, for a total dose of 69 e^-^/Å^2^/s per micrograph. Each H2R/Nb6M/NabFab/anti-Fab Nb image stack is composed of 68 frames with an incident electron dose of 0.86 e^-^/Å^2^ per frame, for a total dose of 58.58 e^-^/Å^2^/s per micrograph.

### Cryo-EM Data Processing

A pictorial depiction of each cryoEM processing workflow can be found in Figs. S2C, S4C, S6C, and S8C. Briefly, dose-fractionated image stacks were imported into RELION-3.1^37^ and subjected to beam-induced motion correction and dose weighting with MotionCor2^38^. Contrast transfer function parameter estimation was performed with CTFFIND-4.1^39^. Particle selection and extraction was performed with the Laplacian autopicker function of RELION-3.1 on micrographs with a CTF fit better than 3.5 Å. The extracted particle stack was imported into CryoSPARC^40^ and subjected to multiple rounds of 2D classification followed by iterative rounds of *ab initio* reconstruction with multiple classes and heterogeneous refinement to further classify particles. In early 3D iterations, particles from poor 3D classes were subjected to additional 2D classification and good particles were ‘rescued’ for further 3D classification. Once iterative *ab initio* and heterogenous refinement no longer produced higher resolution reconstruction, particles were re-imported to RELION-3.1. In the cases of hNTSR1 and MOR CTF fit cutoffs of 3.1 Å and 3.3 Å were applied, respectively. Particles were then subjected to 3D refinement in RELION-3.1 followed by Bayesian polishing. Polished particles were then imported back to CryoSPARC for non-uniform refinement, global CTF refinement based on optics group, and repeated non-uniform and local refinement.

### Model Building and Refinement

All initial models were rigid body fit into cryo-EM maps using the ChimeraX^41^ software. For hNTSR1, the cryo-EM model of active state hNTSR1 was used as the initial model (PDB:6OS9^19^); for MOR the crystal structure of inactive MOR bound to a covalent antagonist was used as the initial model (PDB:4DKL^42^), for SSTR2 a homology model generated from KOR (PDB:6VI4^4^) was used; and for H2R a homology model generated from H1R (PDB:3RZE^33^) was used. In all cases KOR/Nb6 complex (PDB:6VI4^4^) was aligned to the receptor to produce the initial model for the nanobody. An initial structure for the NabFab was taken from PDB:7PHP. All structures were manually refined in COOT^43^ with iterative real-space refinement in Phenix^44^. Once accurate modeling was achieved for the protein components, the GemSpot pipeline^45^ was used for automatic modeling of the ligands alvimopan, SR48692, and famotidine into MOR, hNTSR1, and H2R, respectively. Final refinement was executed in Phenix, followed by model-free Phenix map modification for NTSR1 and MOR^46^. All maps displayed in this work are those prior to model-free map modification unless stated otherwise.

### Molecular Dynamics Simulations

Simulations of the KOR/Nb6 complex started with the crystal structure PDB:6VI4, with the pose of JDTic replaced with that of PDB:4DJH^21^. Maestro’s protein preparation tool was used to assign protonation states, optimize hydrogen bonding, and build missing sidechains and loops. Once the receptor was prepared, the orientation of proteins in membranes webserver (OPM)^47^ was used to align the receptor as it would be in a membrane in the xy plane and the CHARMM-GUI^48^ was used to build the full system with a 1-palmitoyl-2-oleoyl-*sn*-glycero-3-phosphocholine POPC/CHS lipid bilayer, TIP3P water^49^, and 150 mM NaCl. PSF and PDB files were generated with the OPLS-AA/M^50,51^ forcefield in VMD^52^ for the system. For Mb6, a homology model was built from PDB:6XVI with the Prime homology modeling tool^53^. This model was solvated in TIP3P water and 150 mM NaCl in VMD for simulation.

Molecular dynamics simulations were carried out in NAMD^54^ with a 2 fs timestep with SHAKE and SETTLE algorithms^55,56^, with a Langevin thermostat and a Nosé-Hoover Langevin piston barostat at 1 atm with a period of 50 fs and decay of 25 fs. Periodic boundary conditions were used with nonbonded interactions smoothed starting at 10 Å to 12 Å with long-range interactions treated with particle mesh Ewald (PME). The system was minimized for 1,500 steps and then slowly heated from 0 to 303.15K in increments of 20K simulating for 0.4 ns at each increment. For the Kappa/Nb6 simulations, all non-hydrogen, non-water, and non-ion atoms were restrained with a 1 kcal/mol/ Å^2^ harmonic restraint during heating and an additional 10 ns of equilibration. 1 kcal/mol/ Å^2^ harmonic restraints were used for an additional 10 ns of equilibration on all non-hydrogen protein atoms followed by a final 10 ns of restrained equilibration with 1 kcal/mol/ Å^2^ harmonic restraint on only Cα atoms. The first 30 ns of unrestrained molecular dynamics were also considered to be equilibration, with an additional 500 ns of simulation performed. All simulations were performed in triplicate. For Mb6, the first 30 ns of unrestrained molecular dynamics were treated as equilibration, and 100 ns of triplicate molecular dynamics simulations were performed.

### Docking

The cryo-EM structure of active MOR (PDB:6DDE)^3^ was prepared in Maestro and carfentanil was docked with Glide XP docking^57^. The cryo-EM structure of inactive H2R was prepared in Maestro, and ranitidine and cimetidine were docked with Glide XP docking^57^. The crystal structure of inactive H1R (PDB:3RZE) was prepared in Maestro and loratadine and cetirizine were docked with Glide XP docking^57^.

## References

1. Robertson, M. J., Meyerowitz, J. G. & Skiniotis, G. Drug discovery in the era of cryo-electron microscopy. Trends in Biochemical Sciences (2021). doi:10.1016/j.tibs.2021.06.008

2. Hauser, A. S., Attwood, M. M., Rask-Andersen, M., Schiöth, H. B. & Gloriam, D. E. Trends in GPCR drug discovery: New agents, targets and indications. Nat. Rev. Drug Discov. (2017). doi:10.1038/nrd.2017.178

3. Koehl, A. et al. Structure of the μ-opioid receptor-Gi protein complex. Nature 558, 547–552 (2018).

4. Che, T. et al. Nanobody-enabled monitoring of kappa opioid receptor states. Nat. Commun. (2020). doi:10.1038/s41467-020-14889-7

5. Gully, D. et al. Biochemical and pharmacological profile of a potent and selective nonpeptide antagonist of the neurotensin receptor. Proc. Natl. Acad. Sci. U. S. A. (1993). doi:10.1073/pnas.90.1.65

6. Uchanski, T. et al. Megabodies expand the nanobody toolkit for protein structure determination by single-particle cryo-EM. Nat. Methods (2021). doi:10.1038/s41592-020-01001-6

7. Zimmerman, D. M. et al. Discovery of a Potent, Peripherally Selective trans-3,4-Dimethyl-4-(3-hydroxyphenyl)piperidine Opioid Antagonist for the Treatment of Gastrointestinal Motility Disorders. J. Med. Chem. (1994). doi:10.1021/jm00041a003

8. Bloch, J., Mukherjee, S., Kowal, J., Filippova, E. V., Niederer, M., Pardon, E., Seyaert, J., Kossiakoff, A. A., Locher, K. P. Development of a universal nanobody-binding Fab module for fiducial-assisted cryo-EM strudies of membrane proteins P.N.A.S. (2021) doi:10.1073/pnas.2115435118

9. Carraway, R. & Leeman, S. E. Characterization of radioimmunoassayable neurotensin in the rat. Its differential distribution in the central nervous system, small intestine, and stomach. J. Biol. Chem. (1976). doi:10.1016/s0021-9258(17)32938-1

10. Christou, N. et al. Neurotensin pathway in digestive cancers and clinical applications: an overview. Cell Death and Disease (2020). doi:10.1038/s41419-020-03245-8

11. Liu, J. et al. neurotensin receptor 1 antagonist SR48692 improves response to carboplatin by enhancing apoptosis and inhibiting drug efflux in ovarian cancer. Clin. Cancer Res. (2017). doi:10.1158/1078-0432.CCR-17-0861

12. Volkow, N. D. & Blanco, C. The changing opioid crisis: development, challenges and opportunities. Molecular Psychiatry (2021). doi:10.1038/s41380-020-0661-4

13. Volkow, N. D. & Collins, F. S. The Role of Science in Addressing the Opioid Crisis. N. Engl. J. Med. (2017). doi:10.1056/nejmsr1706626

14. Günther, T. et al. International union of basic and clinical pharmacology. CV. somatostatin receptors: Structure, function, ligands, and new nomenclature. Pharmacol. Rev. (2018). doi:10.1124/pr.117.015388

15. Ballesteros, J. A. & Weinstein, H. Integrated methods for the construction of three-dimensional models and computational probing of structure-function relations in G protein-coupled receptors. Methods Neurosci. (1995). doi:10.1016/S1043-9471(05)80049-7

16. Stoeber, M. et al. A Genetically Encoded Biosensor Reveals Location Bias of Opioid Drug Action. Neuron (2018). doi:10.1016/j.neuron.2018.04.021

17. Deluigi, M. et al. Complexes of the neurotensin receptor 1 with small-molecule ligands reveal structural determinants of full, partial, and inverse agonism. Sci. Adv. (2021). doi:10.1126/sciadv.abe5504

18. Mittl, P. R., Ernst, P. & Plückthun, A. Chaperone-assisted structure elucidation with DARPins. Current Opinion in Structural Biology (2020). doi:10.1016/j.sbi.2019.12.009

19. Scott, D. J., Kummer, L., Egloff, P., Bathgate, R. A. D. & Plückthun, A. Improving the apo-state detergent stability of NTS1 with CHESS for pharmacological and structural studies. Biochim. Biophys. Acta - Biomembr. (2014). doi:10.1016/j.bbamem.2014.07.015

20. Kato, H. E. et al. Conformational transitions of a neurotensin receptor 1–Gi1 complex. Nature 572, 80–85 (2019).

21. Russo, C. J. & Passmore, L. A. Ultrastable gold substrates for electron cryomicroscopy. Science (80-.). (2014). doi:10.1126/science.1259530

22. Wu, H. et al. Structure of the human κ-opioid receptor in complex with JDTic. Nature (2012). doi:10.1038/nature10939

23. Van Bever, W. F. M., Niemegeers, C. J. E. & Schellekens, K. H. L. P. A. J. N 4 substituted 1 (2 arylethyl) 4 piperidinyl N phenylpropanamides, a novel series of extremely potent analgesics with unusually high safety margin. Arzneimittel-Forschung/Drug Res. (1976).

24. Robertson, M. J., Meyerowitz, J. M., Panova, O., Borrelli, K. W., Skiniotis, G. Plasticity in Ligand Recognition at Somatostatin Receptors. NSMB (2022)

25. Tunyasuvunakool, K. et al. Highly accurate protein structure prediction for the human proteome. Nature (2021). doi:10.1038/s41586-021-03828-1

26. Baek, M. et al. Accurate prediction of protein structures and interactions using a three-track neural network. Science (80-.). (2021). doi:10.1126/science.abj8754

27. Kong, H., Raynor, K., Yasuda, K., Bell, G. I. & Reisine, T. Mutation of an aspartate at residue 89 in somatostatin receptor subtype 2 prevents Na+ regulation of agonist binding but does not alter receptor-G protein association. Mol. Pharmacol. (1993).

28. Martin, S., Botto, J. M., Vincent, J. P. & Mazella, J. Pivotal role of an aspartate residue in sodium sensitivity and coupling to G proteins of neurotensin receptors. Mol. Pharmacol. (1999). doi:10.1124/mol.55.2.210

29. Yabaluri, N. & Medzihradsky, F. Regulation of μ-opioid receptor in neural cells by extracellular sodium. J. Neurochem. (1997). doi:10.1046/j.1471-4159.1997.68031053.x

30. Fenalti, G. DRUG DISCOVERY Molecular control of delta -opioid receptor signalling. Nat. -LONDON-(2014).

31. Ereño-Orbea J., Sicard T., Cui H., Carson J., Hermans P., Julien J.P. Structural Basis of Enhanced Crystallizability Induced by a Molecular Chaperone for Antibody Antigen-Binding Fragments J. Mol. Biol. (2018)

32. Xu, P. et al. Structural insights into the lipid and ligand regulation of serotonin receptors. Nature (2021) doi:10.1038/s41586-021-03376-8.

33. Shimamura, I., et al. Structure of the human histamine H1 receptor complex with doxepin. Nature (2011) doi:10.1038/nature10246.

34. Marosi, A., Szalay, Z., Béni, S., Szakács, Z., Gáti, T., Rácz, Á., Noszál, B., Demeter, Á. Solution-state NMR spectroscopy of famotidine revisited: spectral assignment, protonation sites, and their structural consequences. Anal. Bioanal. Chem. (2021) doi:10.1007/s00216-011-5599-6.

35. Gantz, I., DelValle, J., Wang. L-D., Tashiro, T., Munzert, G., Guo, Y-J., Konda, Y., Yamada, Y. Molecular Basis for the Interaction of Histamine with the Histamine H2 Receptor. J. Biol. Chem. (1992) doi:10.1016/S0021-9258(19)36764-X.

36. Towns, J. et al. XSEDE: Accelerating scientific discovery. Comput. Sci. Eng. (2014). doi:10.1109/MCSE.2014.80

37. Zivanov, J. et al. RELION-3: New tools for automated high-resolution cryo-EM structure determination. bioRxiv (2018). doi:10.1101/421123

38. Zheng, S. Q. et al. MotionCor2: Anisotropic correction of beam-induced motion for improved cryo-electron microscopy. Nature Methods 14, 331–332 (2017).

39. Rohou, A. & Grigorieff, N. CTFFIND4: Fast and accurate defocus estimation from electron micrographs. J. Struct. Biol. 192, 216–221 (2015).

40. Punjani, A., Rubinstein, J. L., Fleet, D. J. & Brubaker, M. A. CryoSPARC: Algorithms for rapid unsupervised cryo-EM structure determination. Nat. Methods (2017). doi:10.1038/nmeth.4169

41. Pettersen, E. F. et al. UCSF ChimeraX: Structure visualization for researchers, educators, and developers. Protein Sci. (2021). doi:10.1002/pro.3943

42. Manglik, A. et al. Crystal structure of the μ-opioid receptor bound to a morphinan antagonist. Nature (2012). doi:10.1038/nature10954

43. Emsley, P., Lohkamp, B., Scott, W. G. & Cowtan, K. Features and development of Coot. Acta Crystallogr. Sect. D Biol. Crystallogr. 66, 486–501 (2010).

44. Adams, P. D. et al. PHENIX: A comprehensive Python-based system for macromolecular structure solution. Acta Crystallogr. Sect. D Biol. Crystallogr. 66, 213–221 (2010).

45. Robertson, M. J., van Zundert, G. C. P., Borrelli, K. & Skiniotis, G. GemSpot: A Pipeline for Robust Modeling of Ligands into Cryo-EM Maps. Structure (2020). doi:10.1016/j.str.2020.04.018

46. Terwilliger, T. C., Ludtke, S. J., Read, R. J., Adams, P. D. & Afonine, P. V. Improvement of cryo-EM maps by density modification. Nat. Methods (2020). doi:10.1038/s41592-020-0914-9

47. Lomize, M. A., Pogozheva, I. D., Joo, H., Mosberg, H. I. & Lomize, A. L. OPM database and PPM web server: Resources for positioning of proteins in membranes. Nucleic Acids Res. 40, (2012).

48. Lee, J. et al. CHARMM-GUI Input Generator for NAMD, GROMACS, AMBER, OpenMM, and CHARMM/OpenMM Simulations Using the CHARMM36 Additive Force Field. J. Chem. Theory Comput. 12, 405–413 (2016).

49. Jorgensen, W. L., Chandrasekhar, J., Madura, J. D., Impey, R. W. & Klein, M. L. Comparison of simple potential functions for simulating liquid water. J. Chem. Phys. 79, 926–935 (1983).

50. Robertson, M. J., Tirado-Rives, J. & Jorgensen, W. L. Improved Peptide and Protein Torsional Energetics with the OPLS-AA Force Field. J. Chem. Theory Comput. 11, 3499– 3509 (2015).

51. Robertson, M. J., Skiniotis, G. Development of OPLS-AA/M Parameters for Simulations of G-Protein Coupled Receptors and other Membrane Proteins. bioRxiv. (2022).

52. Humphrey, W., Dalke, A. & Schulten, K. VMD: Visual molecular dynamics. J. Mol. Graph. 14, 33–38 (1996).

53. Jacobson, M. P. et al. A Hierarchical Approach to All-Atom Protein Loop Prediction. Proteins Struct. Funct. Genet. (2004). doi:10.1002/prot.10613

54. Phillips, J. C. et al. Scalable molecular dynamics with NAMD. Journal of Computational Chemistry 26, 1781–1802 (2005).

55. Ryckaert, J. P., Ciccotti, G. & Berendsen, H. J. C. Numerical integration of the cartesian equations of motion of a system with constraints: molecular dynamics of n-alkanes. J. Comput. Phys. (1977). doi:10.1016/0021-9991(77)90098-5

56. Miyamoto, S. & Kollman, P. A. Settle: An analytical version of the SHAKE and RATTLE algorithm for rigid water models. J. Comput. Chem. (1992). doi:10.1002/jcc.540130805

57. Friesner, R. A. et al. Extra precision glide: Docking and scoring incorporating a model of hydrophobic enclosure for protein-ligand complexes. J. Med. Chem. 49, 6177–6196 (2006).

