## Supplementary material for "Structure Determination of Inactive-State GPCRs with a Universal Nanobody": All supplemental files

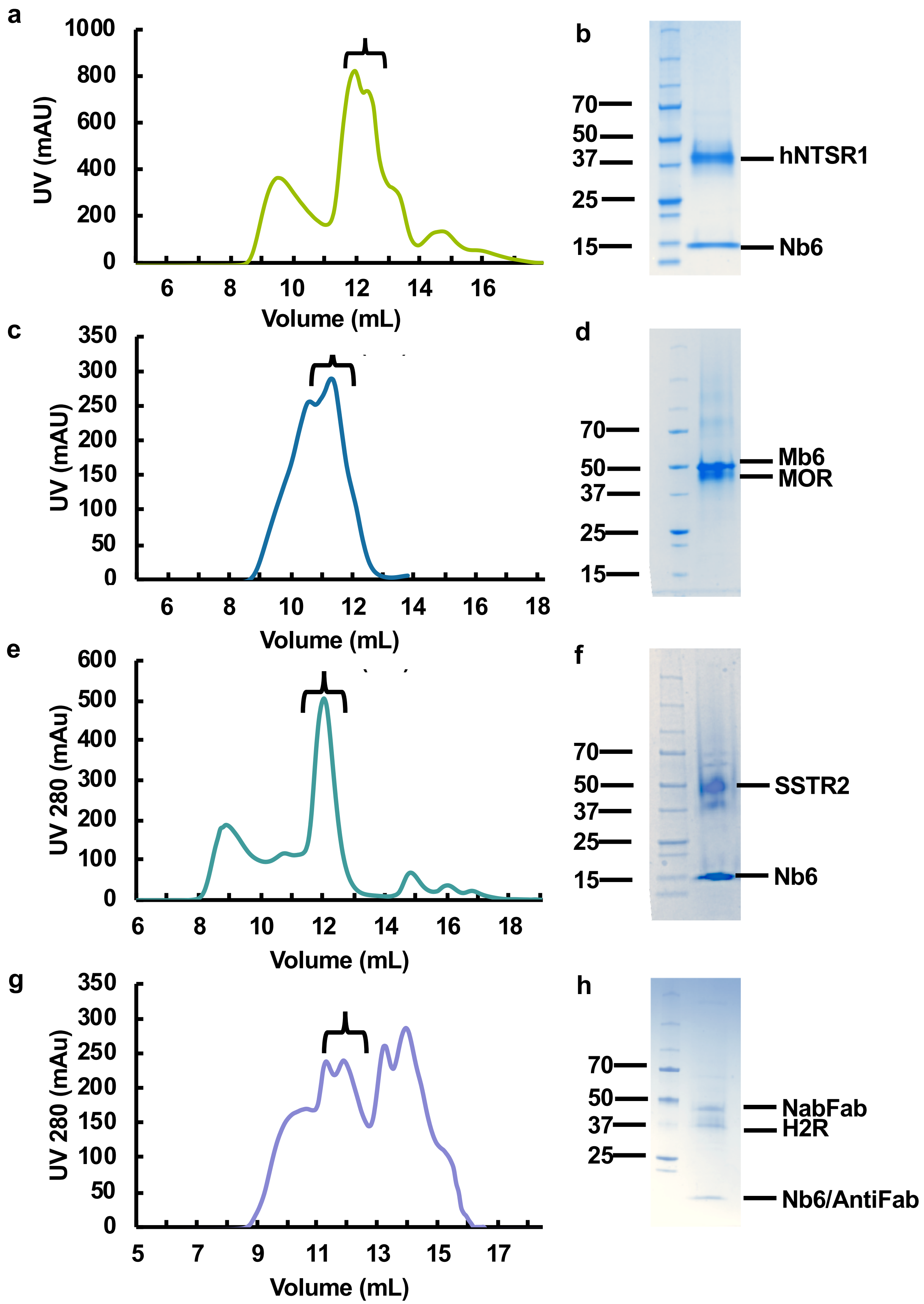


**Extended Data Fig. 1 | Biochemical characterization of receptor Nb6/Mb6 complexes. a,** Size exclusion chromatography (SEC) profile of hNTSR1/Nb6 complex; bracket indicates fractions harvested for cryoEM. **b,** SDS-PAGE gel of hNTSR1/Nb6 complex. **c,** SEC profile of MOR/Mb6 complex; bracket indicates fractions harvested for cryoEM. **d,** SDS-PAGE gel of MOR/Mb6 complex. **e,** SEC profile of SSTR2/Nb6 complex; bracket indicates fractions harvested for cryoEM. **f,** SDS-PAGE gel of SSTR2/Nb6 complex. **g,** SEC profile of H2R/Nb6M/NabFab/Anti-Fab Nb **h,** SDS-PAGE gel of H2R/Nb6M/NabFab/Anti-Fab Nb

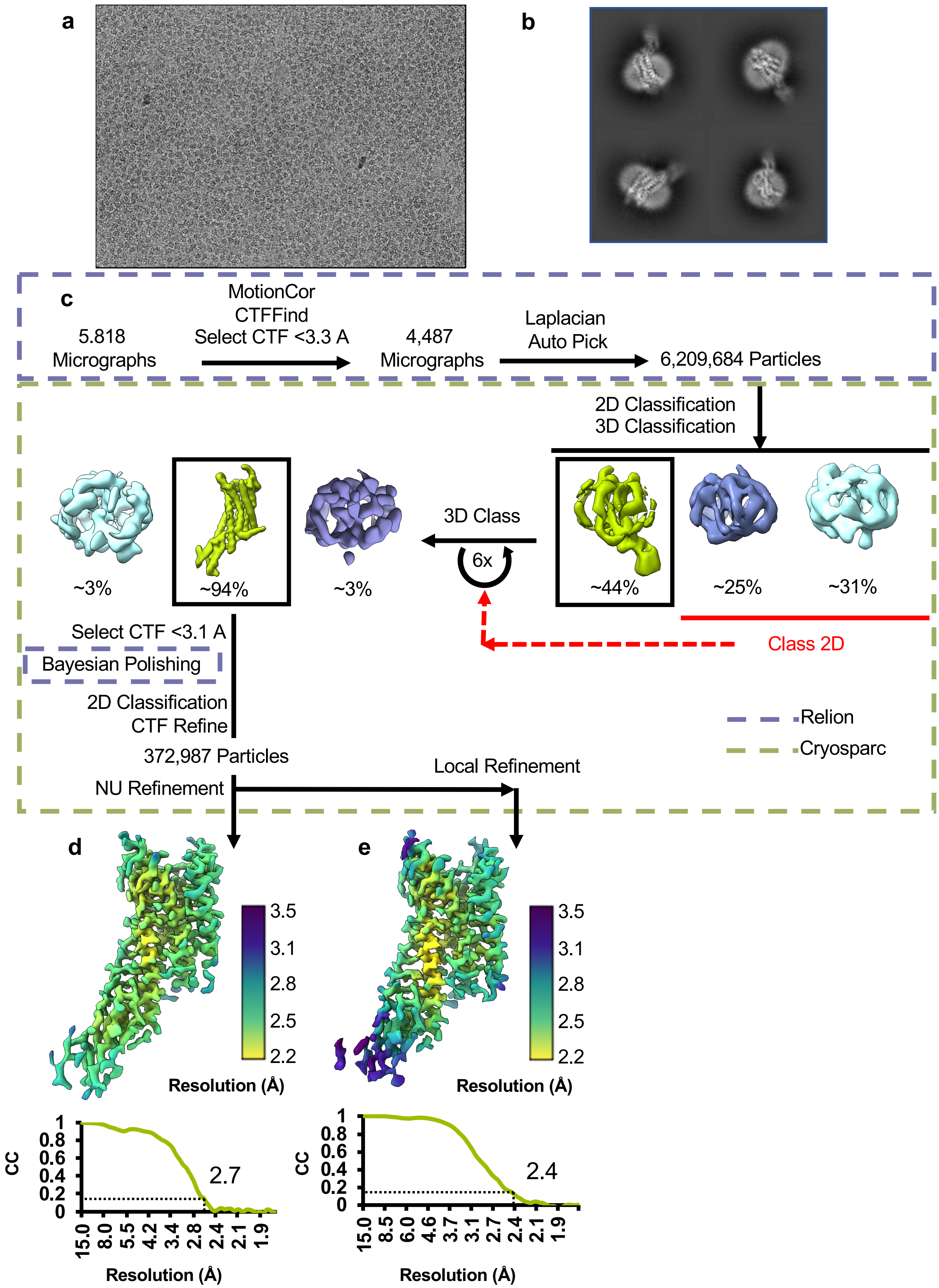


**Extended Data Fig. 2 | hNTSR1/Nb6 cryo-EM data collection and processing. a,** Representative micrograph of hNTSR1/Nb6 complex. **b,** Example final 2D classes of hNTSR1/Nb6 complex. **c,** Cryo-EM data processing workflow. **d,** Local resolution of hNTSR1/Nb6 global refinement with FSC curve below. **e,** Local resolution of hNTSR1/Nb6 local refinement with FSC curve below.

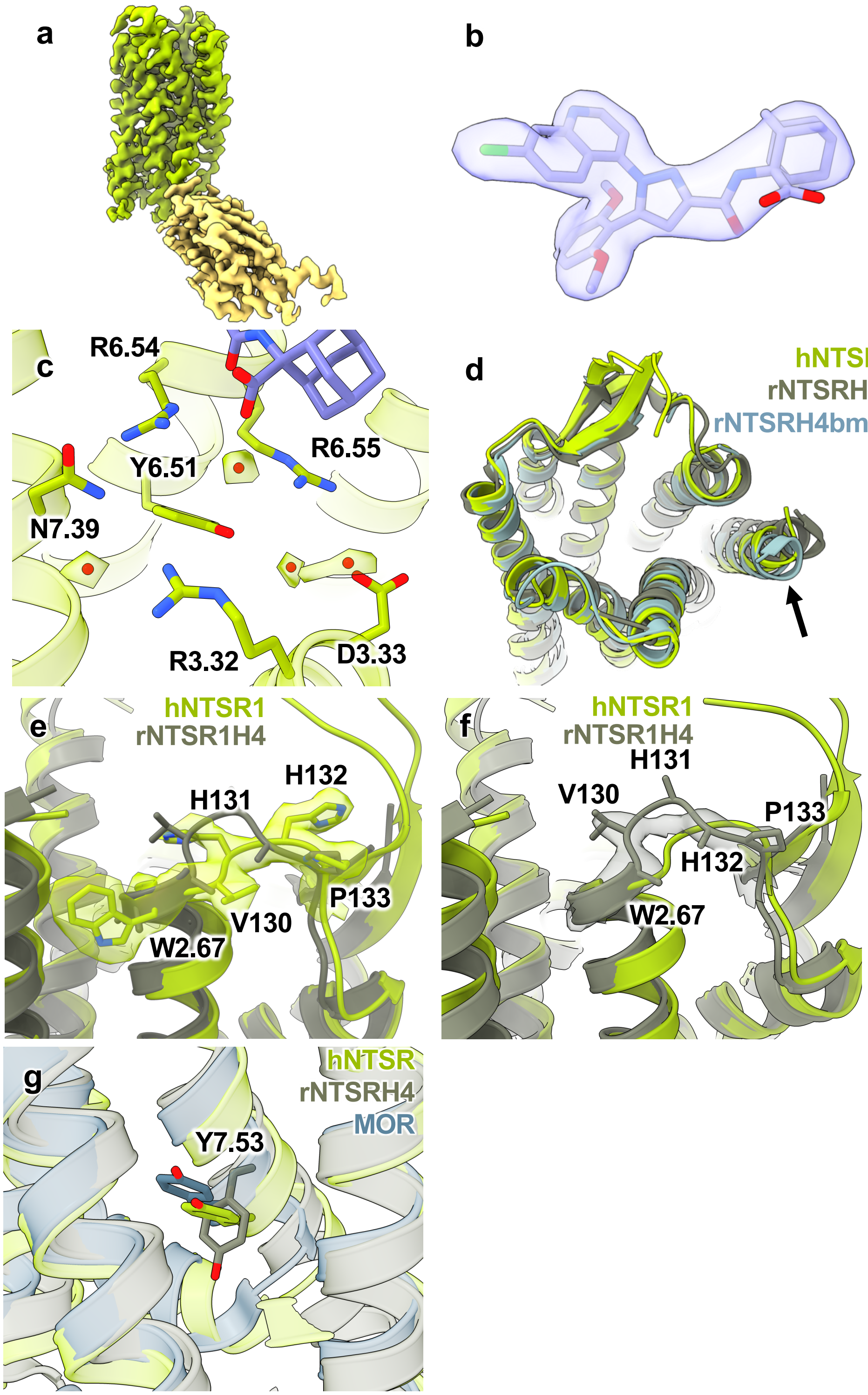


**Extended Data Fig. 3 | Comparison between hNTSR1/Nb6 cryoEM structure and rNTSR1H4/DARPin crystal structure. a,** 2Fo-Fc crystallography map at 2.7 Å of rNTSR-H4bmx contoured at $\sigma$=1.0. **b,** 2Fo-Fc crystallography map at 2.7 Å of rNTSR-H4bmx bound SR antagonist contoured at $\sigma$=1.0. **c,** Structure of hNTSR1 (green) in the extracellular ligand binding pocket showing water molecules and corresponding cryoEM map features. **d,** Overlay of hNTSR1 (green) and rNTSR1H4 (gray) highlighting the movement of TM1 **e,** Overlay of hNTSR1 (green) and rNTSR1H4 (gray) with the cryoEM map for hNTSR1 (green) for ECL1 **f,** Overlay of hNTSR1 (green) and rNTSR1H4 (gray) with the rNTSR1H4 crystal structure 2Fo-Fc map contoured at $\sigma$=1.0. **g,** Overlay of hNTSR1 (green), rNTSR1H4 (gray), and MOR PDB:4DKL (blue) showing the position of NPXXY motif Y7.53.

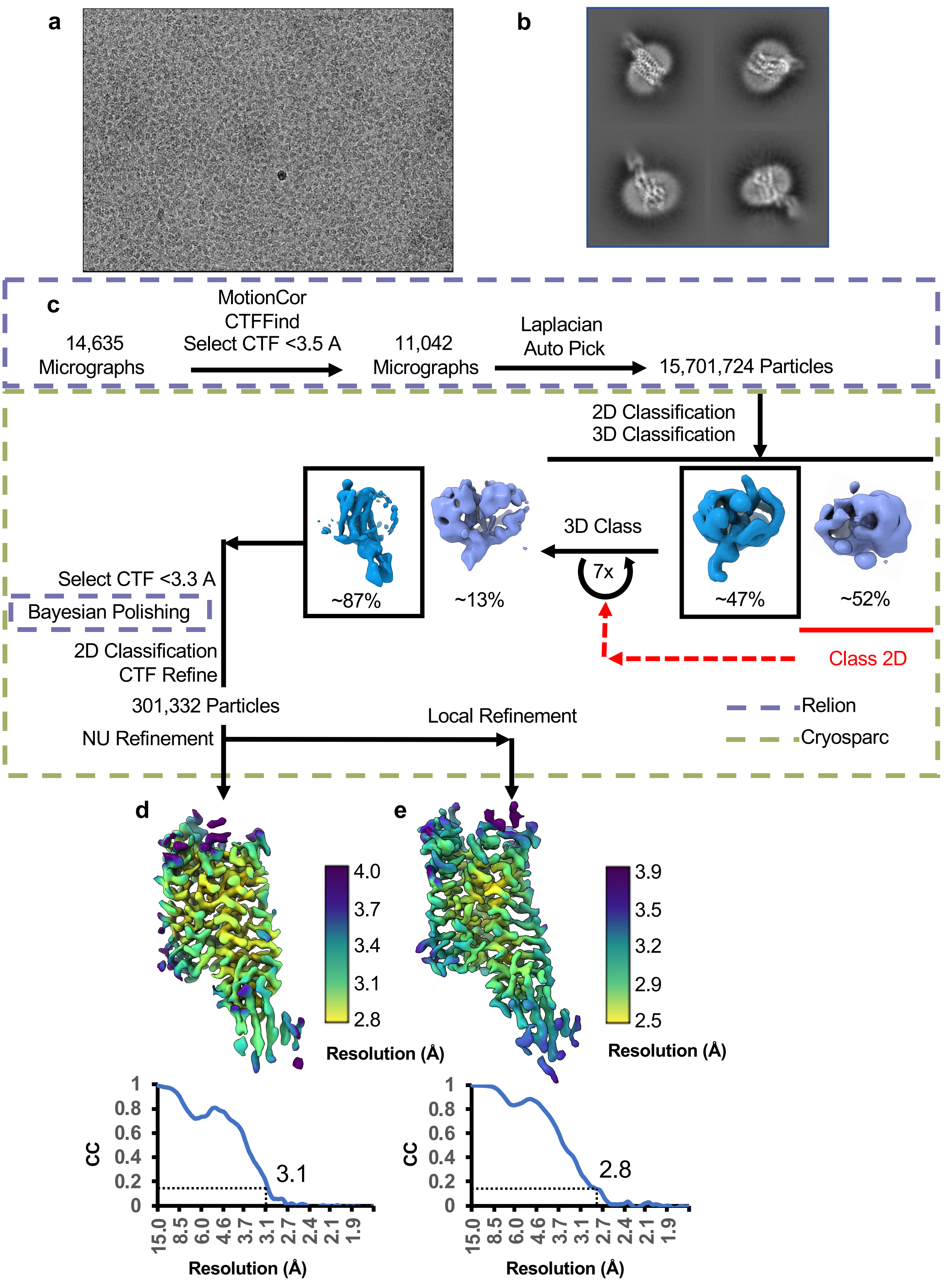


**Extended Data Fig. 4 | MOR/Mb6 cryo-EM data collection and processing. a,** Representative micrograph of MOR/Mb6 complex. **b,** Example final 2D classes of MOR/Mb6 complex. **c,** Cryo-EM data processing workflow. **d,** Local resolution of MOR/Mb6 global refinement with FSC curve below. **e,** Local resolution of MOR/Mb6 local refinement with FSC curve below.

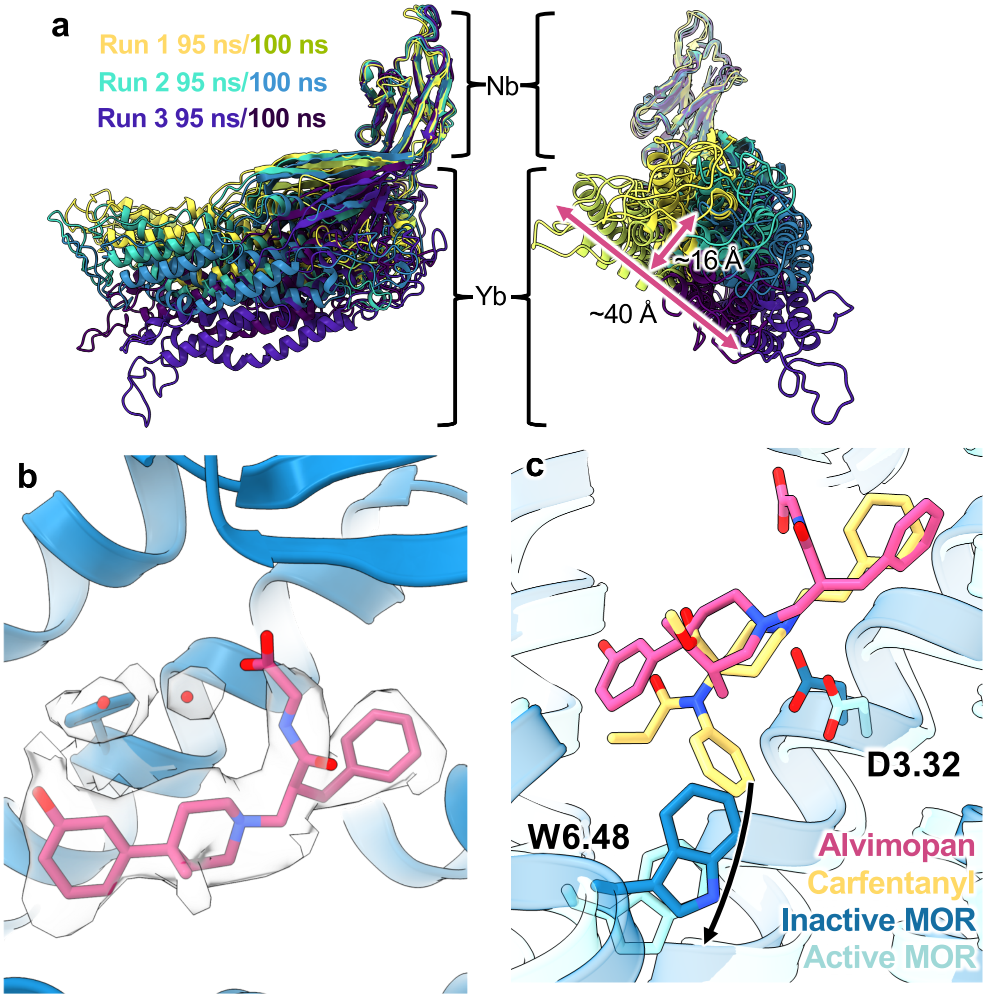


**Extended Data Fig. 5 | Inactive MOR bound to avlimopan. a,** overlay of snapshots from Mb6 molecular dynamics simulations aligned on the nanobody portion. **b,** Comparison of Inactive MOR (dark blue) bound to alvimopan (magenta) and bound water in density modified map (grey) **c,** Comparison of Inactive MOR (dark blue) bound to alvimopan (magenta) and carfentanyl (yellow) docked into the active state MOR receptor (light blue).

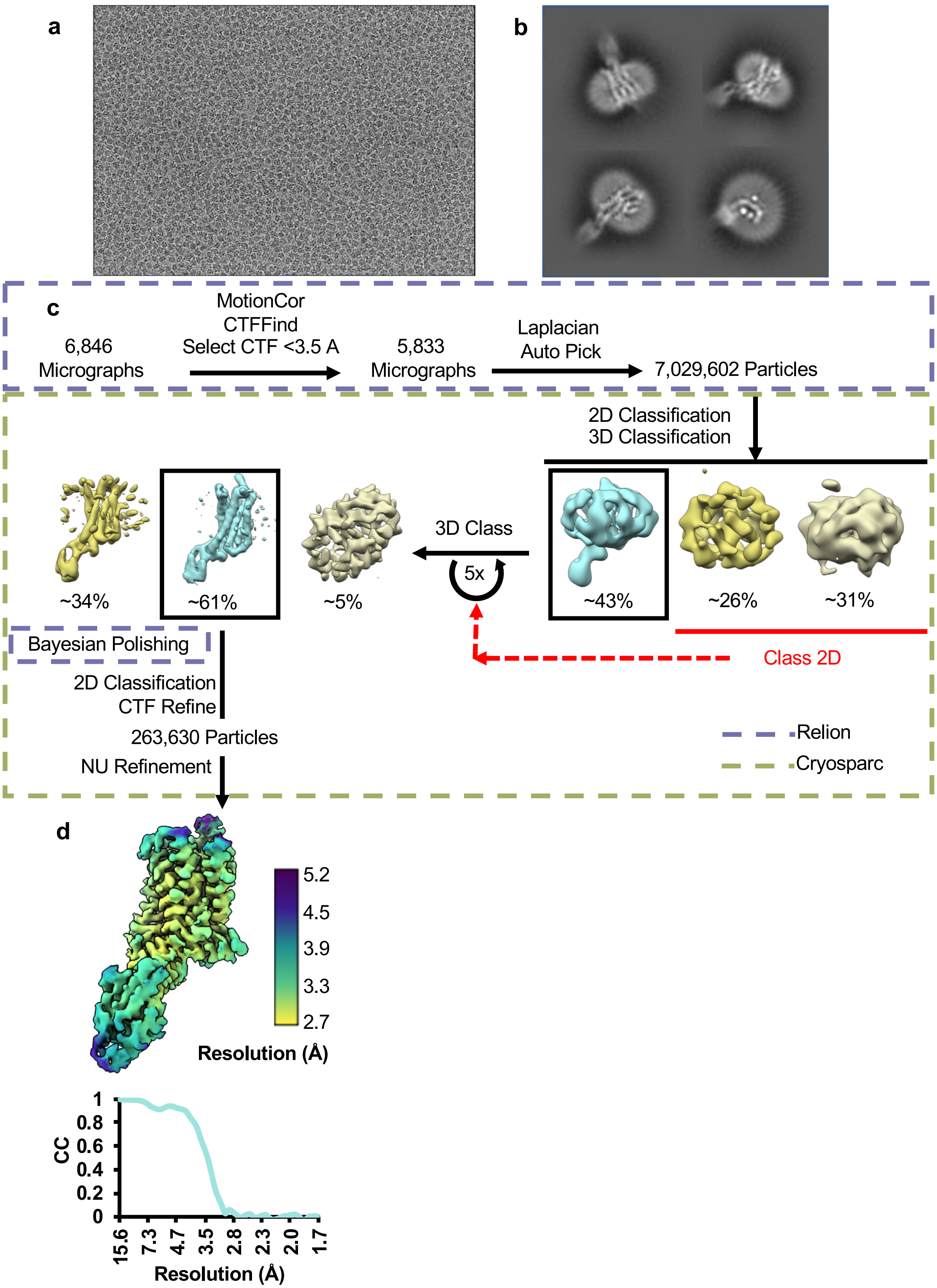


**Extended Data Fig. 6 | SSTR2/Nb6 cryo-EM data collection and processing. a,** Representative micrograph of SSTR2/Nb6 complex. **b,** Example final 2D classes of SSTR2/Nb6 complex. **c,** Cryo-EM data processing workflow. **d,** Local resolution of SSTR2/Nb6 global refinement with FSC curve below.

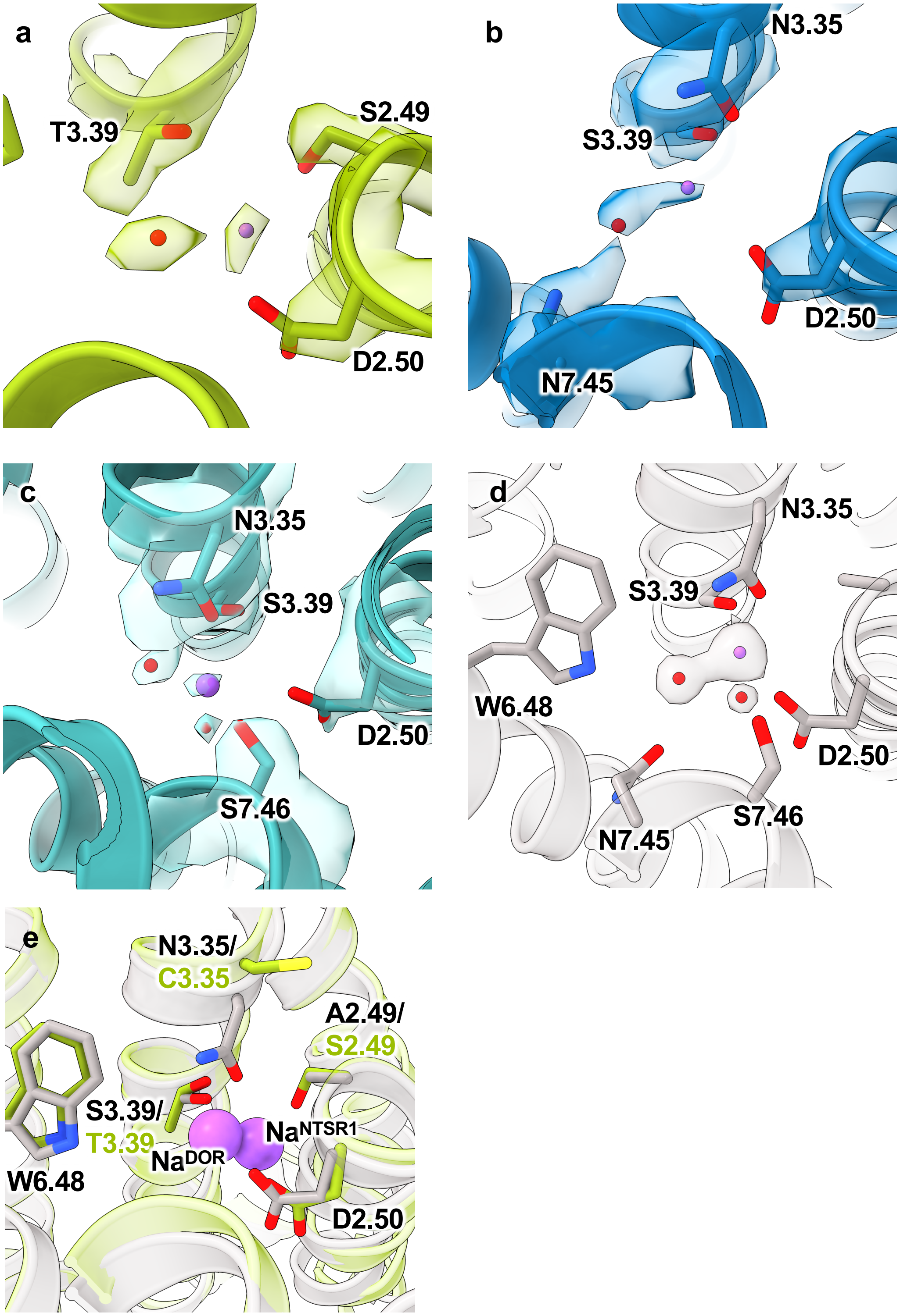


**Supplemental Fig. 7 | Comparison of CryoEM Maps and Putative Ion Sites. a.** Sodium ion site of hNTSR1 with cryoEM map. **b,** Probable sodium ion site of MOR with cryoEM map. **c,** Probable sodium ion site of SSTR2 with cryoEM map. **d,** Sodium ion site of DOR (PDB:4N6H) with 2Fo-Fc map contoured at $\sigma$=2.0 **e,** Overlay of NTSR1 (green) cryoEM structures sodium ion binding site with the DOR sodium coordination site structure (gray, PDB:4N6H).

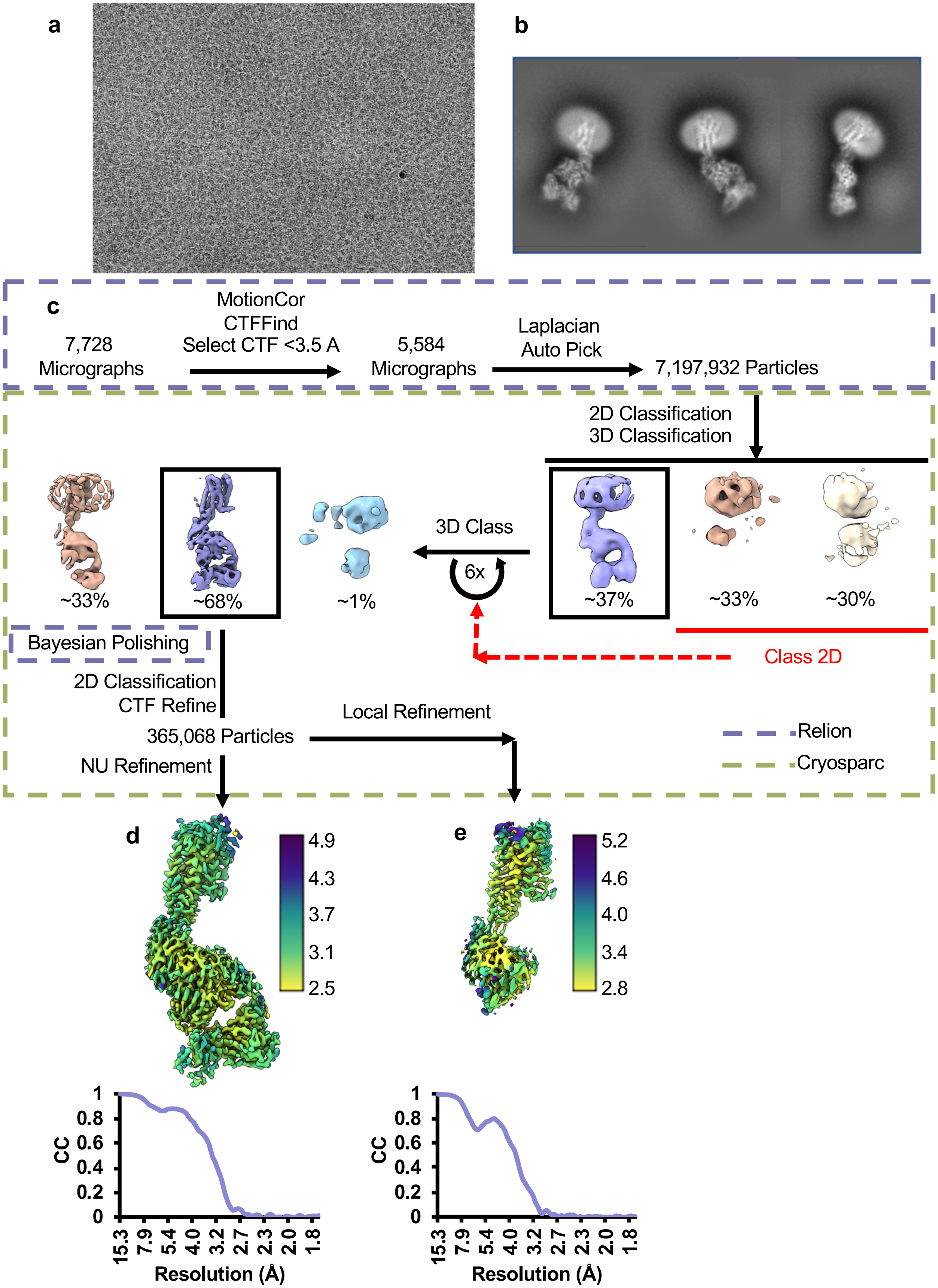


**Extended Data Fig. 8 | Inactive H2R/Nb6M/NabFab/Anti-Fab Nb cryoEM data collection and processing. a**, Representative micrograph of H2R complex. **b,** Example final 2D classes of H2R complex. **c,** CryoEM data processing workflow. **d,** Local resolution of H2R global refinement **e,** Local resolution of H2R local refinement **f,** FSC curve of H2R global refinement G) FSC curve of H2R local refinement

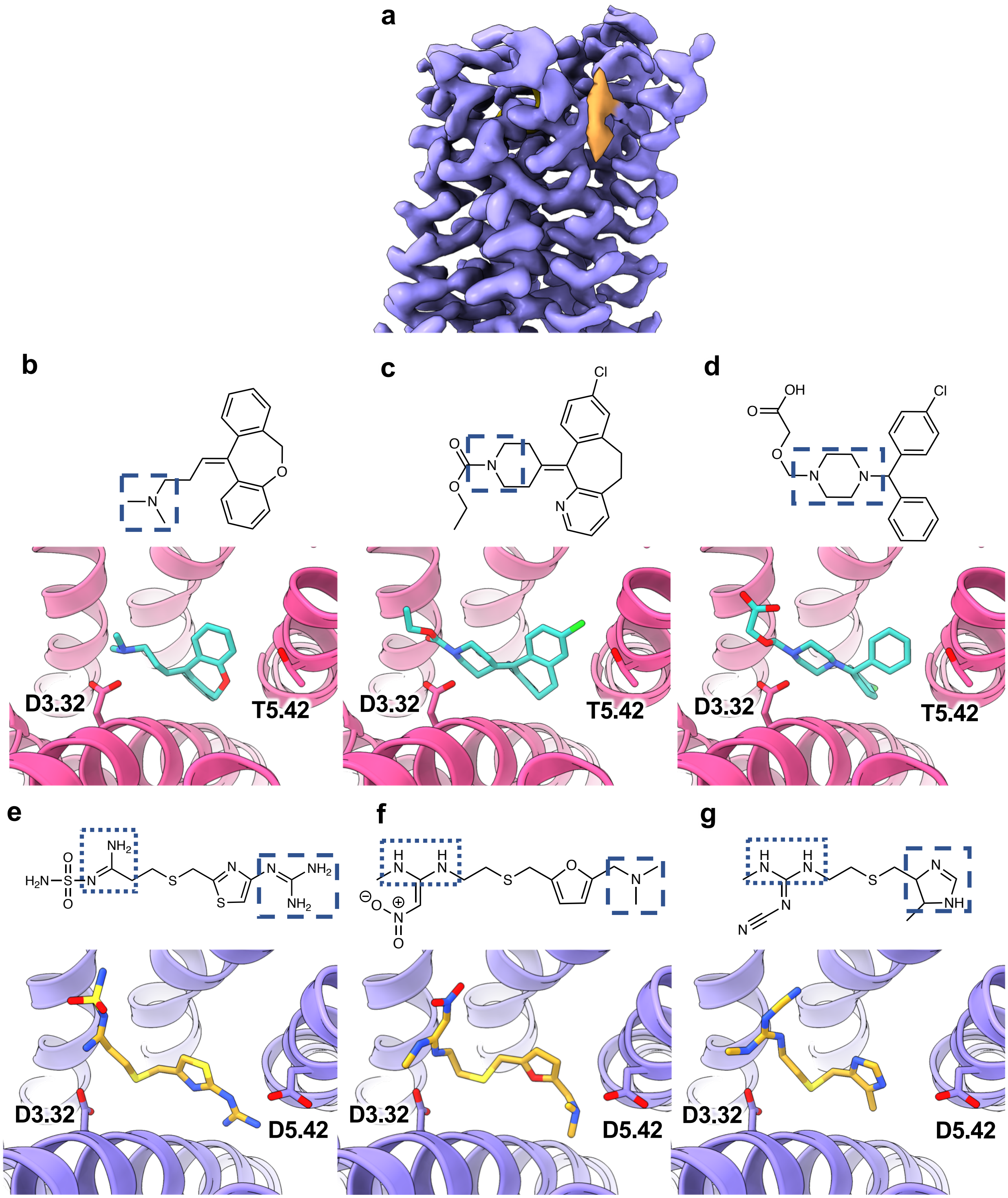


**Extended Data Fig. 9 | Inactive H2R Structure and Comparison to H1R a,** Cryo-EM map of H2R with lipid density between TM1 and TM7 colored in orange. **b-d,** Chemical structures of doxepin **b**, loratadine **c**, and cetirizine **d** with protonated amine highlighted and crystal structure of H1R (magenta) bound to doxepin (teal) (PDB:3RZE) loratadine (teal, docked pose) and cetirizine (teal, docked pose). **e-g,** Chemical structures of famotidine **e**, ranitidine **f**, and cimetidine **g** with protonated amines highlighted in dashed (high pKa) and dotted (low pKa) boxes, together with cryoEM structure of H2R (lavender) bound to famotidine (goldenrod, this work), ranitidine (goldenrod, docked pose) and cimetidine (goldenrod, docked pose).

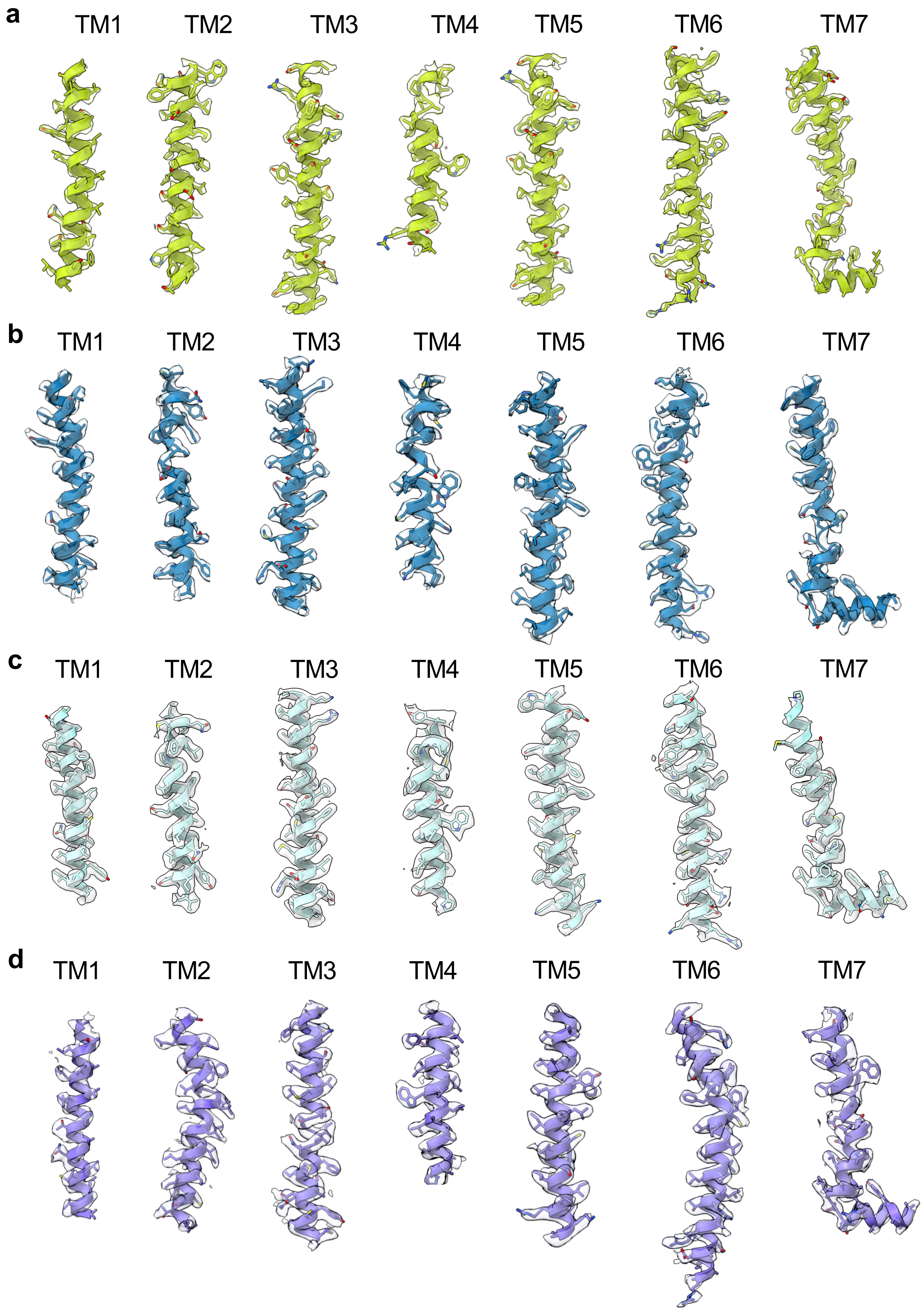


**Extended Data Fig. 10 | Map-Model Agreements. a,** Map-model comparison for NTSR1. **b,** Map-model comparison for MOR. **c,** Map-Model comparison for SSTR2. **d,** Map-Model comparison for H2R.

**Supplemental Table 1. Cryo-EM data collection, refinement and validation statistics**

| | SSTR2/Nb6 | hNTSR1/Nb6 | MOR/Mb6 | H2R/Nb6M/Nab Fab |
|---|---|---|---|---|
| **Data collection and processing** | | | | |
| Voltage (kV) | 300 | 300 | 300 | 300 |
| Electron exposure (e–/Å2) | 68.93 | 60.82 | 61.38 | 58.58 |
| Defocus range (μm) | -0.8 to -1.8 | -0.5 to -1.5 | -0.7 to -1.5 | -0.6 to -1.2 |
| Pixel size (Å) | 0.8521 | 0.8677 | 0.8677 | 0.8677 |
| Symmetry imposed | C1 | C1 | C1 | C1 |
| Initial particle images (no.) | 7,029,602 | 6,209,684 | 15,701,724 | 7,197,932 |
| Final particle images (no.) | 263,630 | 372,987 | 301,332 | 365,068 |
| Map resolution (Å) | 3.1 | 2.8 | 2.4 | 3.0 |
| FSC threshold | 0.143 | 0.143 | 0.143 | 0.143 |
| Map sharpening B factor ($Å^2$) | | | | |
| **Refinement** | | | | |
| Initial Model Used (PDB Code) | 6VI4 | 6OS9, 6VI4 | 4DKL, 6VI4 | 7PHP |
| Model Resolution | 3.0 | 2.4 | 2.8 | 2.9 |
| FSC Threshold | 0.143 | 0.143 | 0.143 | 0.143 |
| *Model Composition* | | | | |
| Non-hydrogen Atoms | 2780 | 2624 | 2556 | 5508 |
| Protein Atoms | | | | |
| Waters & Ions | | | | |
| *B factor ($Å^2$)* | | | | |
| Protein Atoms | 49.06 | 15.27 | 19.41 | 41.22 |
| Waters & Ions | | 6.38 | 10.25 | |
| *R.M.S Deviations* | | | | |
| Bonds (Å) | 0.004 | 0.007 | 0.006 | 0.004 |
| Angles (º) | 0.626 | 0.996 | 0.940 | 0.713 |
| *Validation* | | | | |
| MolProbity score | 1.31 | 1.36 | 1.11 | 1.49 |
| Clashscore | 3.67 | 6.58 | 3.12 | 5.82 |
| Poor rotamers (%) | 0.00 | 0.84 | 0.00 | 0.00 |
| *Ramachandran Plot* | | | | |
| Favored (%) | 97.18 | 98.48 | 97.96 | 97.01 |
| Allowed (%) | 2.82 | 1.52 | 2.04 | 2.99 |
| Outliers (%) | 0.0 | 0.00 | 0.00 | 0.00 |
